# Metabolite-sensitive cross-bridge models of human atria reveal the impact of diabetes on muscle mechanoenergetics

**DOI:** 10.1101/2025.02.28.640209

**Authors:** JH Musgrave, J-C Han, M-L Ward, AJ Taberner, K Tran

## Abstract

Type 2 diabetes is associated with a range of adverse health outcomes, including metabolic dysfunction and increased risk of heart failure. Although the interactions between diabetes and heart disease are complex and incompletely understood, they can be more effectively investigated using multiscale, biophysically-based models of the underlying physiological processes. In this study, experimental data from non-diabetic and diabetic human atrial muscles were used to develop metabolite-sensitive cross-bridge models representing each group. The parameterisation of these cross-bridge models revealed that reduced muscle stress development and a leftward shift of the complex modulus measured in the diabetic muscles could be attributed to reduced cross-bridge stiffness and slower cross-bridge detachment rates, respectively. These cross-bridge models were also integrated into muscle models to investigate the effects of diabetic cross-bridge function, Ca^2+^ handling and altered metabolite availability on isometric and physiological work-loop contractions. The diabetic model produced isometric twitches with lower amplitude and prolonged duration and, in response to lowered ATP concentration, the diastolic stress increased notably. In work-loop simulations, the diabetic model exhibited slower shortening, reduced work output and lower power of shortening. However, it was also more efficient and had a less pronounced negative response to increases in P_i_ concentration. These simulations demonstrate that while experimentally measured differences in diabetic cardiac tissues can lead to impaired function during physiological contractions, they may also offer compensatory advantages. The insights of this study offer clear mechanisms of mechanoenergetic dysfunction in diabetic heart muscle, identifying potential therapeutic targets to improve cardiac outcomes for individuals with diabetes.

## 1 Introduction

As the prevalence of type 2 diabetes continues to rise [1], it has become increasingly important to understand the cardiovascular complications associated with this disease. Individuals with type 2 diabetes face a risk of developing heart failure that is at least double that of the general population [2]. Heart failure, a condition in which the heart cannot pump enough blood to meet the body’s demands, is associated with a four-fold increase in mortality risk among diabetic patients [3]. The effects of diabetes on the heart occur across mechanical and energetic domains and span sub-cellular to organ-level scales [2, 4]. Given the complexity of these interactions, much remains unknown about the relationship between diabetes and the development of cardiomyopathy.

Cardiac mechanoenergetics encompasses the study of the processes associated with cardiac contraction and the corresponding expenditure of energy required to sustain it. As a metabolic disease, diabetes is known to affect mitochondrial function in the heart [4, 5]. The downstream effects of this metabolic stress and impairment are varied, ranging from altered availability of energy substrates within the heart [6, 7] to action potential abnormalities [8] and alterations in the sensitivity of myofilaments to Ca^2+^ [9]. Given the critical link between energy metabolism and mechanical function, these impairments ultimately impact the core mechanical function of the heart. At the cellular and tissue levels, the most commonly observed effects are prolonged twitch duration [10, 11] and reduced stress production [9, 12]. A previous study on type 1 diabetic rat heart tissues found no significant changes in work output, heat production or efficiency [10]. However, comparable mechanoenergetics data for human cardiac tissues are lacking, leaving a gap in our understanding of these processes in human physiology. In the absence of such data, mathematical modelling offers a way to bridge this gap by providing insights into the underlying mechanisms and generating testable hypotheses.

The application of mathematical modelling techniques to the fields of physiology and medicine offers insights beyond what can be achieved solely with experimental measurement in two key ways. Firstly, biophysically-based models parameterised using experimental data enable a deeper and more detailed exploration into the mechanisms driving observed behaviour. Secondly, these models can predict responses beyond the specific conditions under which the data were collected. For example, they allow for the testing of hypotheses that cannot be easily examined experimentally and allow for the prediction of interactions when integrated with other biophysical processes.

In a previous study, we performed a comprehensive experimental analysis of cross-bridge function and its responsiveness to metabolites in cardiac muscle isolated from the atria of individuals with diabetes [13]. These experiments revealed three key differences in the diabetic muscles: lower stress production, a leftward shift in the complex modulus - which is indicative of slower cross-bridge cycling - and altered metabolite sensitivity. The development of a mathematical model that captures these experimental effects would offer insights into diabetic heart disease that extend beyond what experiments alone can provide. Specifically, a biophysical metabolite-sensitive cross-bridge model can identify the mechanisms underlying dysfunction in diabetic cross-bridges and predict how these dysfunctions interact with other processes involved in *in vivo* cardiac function.

This study thus describes the development of models of non-diabetic and diabetic human atrial muscle. Leveraging the data from Musgrave et al. [13], we parameterise metabolite-sensitive cross-bridge models using methods similar to those applied in our previous study on rat [14]. The cross-bridge models are then integrated into whole muscle models to simulate Ca^2+^-activated dynamic contractions in the non-diabetic and diabetic muscles. These model simulations allow for the identification of modes and mechanisms of mechanoenergetic impairment in diabetic heart tissue and may be used to assess potential therapeutic pathways.

## 2 Methods

### 2.1 Experimental data

The human atrial models developed in this study were parameterised using mechanical measurements from non-diabetic and diabetic human atrial trabeculae, as detailed in Musgrave et al. [13]. In brief, these data were collected from 10 trabeculae per patient group, sourced from right atrial appendage samples gathered during coronary artery bypass graft surgery. Trabeculae were permeabilised prior to the experiments so that measurements characterising cross-bridge properties could be taken under maximal Ca^2+^ activation (pCa 4.5) at a range of different concentrations of ATP and P_i_. Passive measurements were also made when the trabeculae were exposed to low concentrations of Ca^2+^ (pCa 9.0).

The primary data set used in model fitting was the active complex modulus - the stiffness measured under sinusoidal length perturbations across a range of frequencies. This measurement was taken at a baseline metabolite condition (5 mM ATP and 1 mM P_i_) as well as at four other conditions with variation of one of these metabolites: 1 mM ATP, 0.1 mM ATP, 0 mM P_i_, 10 mM P_i_. The maximum stress (force divided by trabecula cross-sectional area) generated at each of these metabolite conditions was also used to guide the cross-bridge model calibration.

The trabeculae from which these mechanical data were collected were also fixed and immunolabelled for structural imaging, allowing for quantification of the proportion of myofilament content within each of the two groups of trabeculae.

Complex modulus and steady-state force-length measurements made under relaxed conditions were used to parameterise the passive components of the muscle model.

The full set of data used to parameterise the models is available at https://doi.org/10.6084/m9.figshare.28180337.v1.

### 2.2 Cross-bridge model

#### 2.2.1 Model description

The systematic fitting approach outlined in Musgrave et al. [14] was applied to determine the optimal combination of strain and metabolite dependencies for describing the human data. Our human data did not show the increased dip magnitude at low ATP concentrations seen in rat data [14], so it was likely that we would need to employ a different combination of these dependencies. For both the non-diabetic and diabetic groups, the data sets describing the average active complex moduli and active stresses across the five metabolite conditions were used in model fitting. As previously, 64 permutations of the linearised cross-bridge model were fitted to these data by minimising an objective function (Eq. 15). Following systematic model fitting, the objective value was found as a percentage of the range of each set of data to produce the normalised RMSE. The final version of the model was selected based on that which provided the lowest normalised RMSE averaged across the two groups.

The final human atrial cross-bridge model is a three-state stiffness-distortion model [15], as depicted in Fig. 1. We are initially concerned only with the myosin cross-bridge component of the model, where we assume that the thin filament is maximally-activated by Ca^2+^, ignoring the non-permissive state, **N**. The active stress (force normalised to cross-sectional area) produced by the cross-bridge model is:

**Figure 1:**
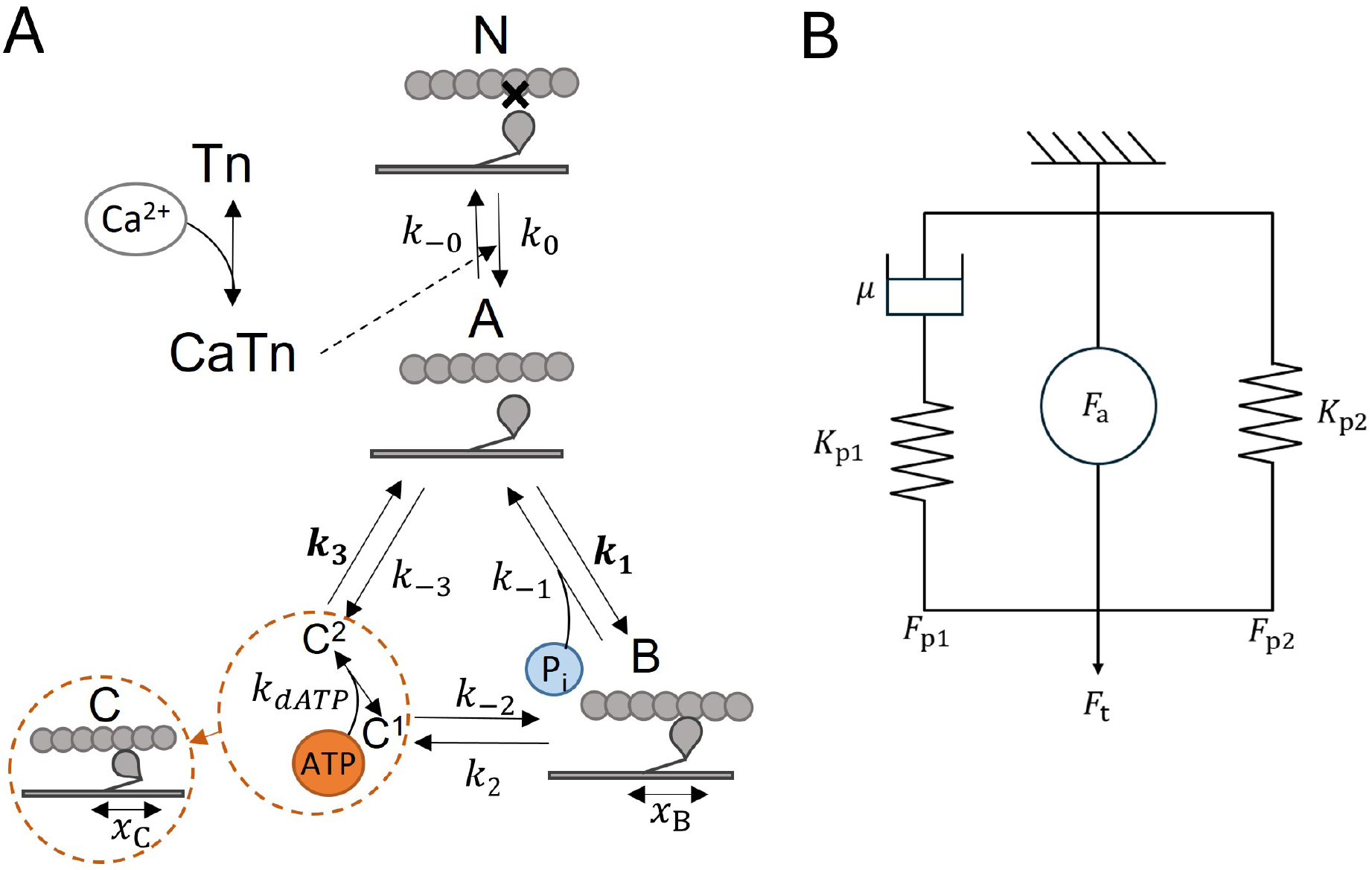
The human muscle model. (A) The cross-bridge model has three main states (States **A, B** and **C**) and is sensitive to P_i_ and ATP. State **C** consists of substates C1 and C2. The bolded transition rates (*k*_*i*_) are dependent on cross-bridge strain. State **N** is a non-permissive state where cross-bridges cannot form. Thin filament regulation by Ca^2+^ allows cross-bridges to transition from State **N** to State **A.** (B) Model of total muscle force consisting of a spring (*K*_p1_) and damper (*µ*) in series, which is in parallel with a non-linear spring (*K*_p2_) and an active force element (*F*_a_) that is the cross-bridge model.

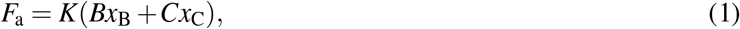

where *F*_a_ is the active stress generated by attached cross-bridges; *B* and *C* are state variables representing the proportion of cross-bridges in each of the two attached states, pre- and post-power stroke, respectively; *x*_B_ and *x*_C_ are state variables representing the mean values of cross-bridge strain in the pre-power stroke state **B** and post-power stroke state **C**, respectively; *K* is a spring constant representing the stiffness of the collective myosin heads. As *K* reflects both the stiffness of an individual myosin head and the number of these for a given cross-section of muscle, and we have measurements reflecting how much of the muscle was occupied by myofilaments, this value can be described more precisely in our model. We thus split *K* into two components:

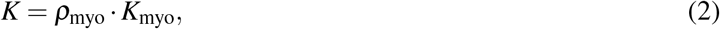

where *ρ*_myo_ is the fraction of the trabecula cross-section made up of myocytes, and *K*_myo_ is the stiffness coefficient of the myocytes within the trabeculae. When the cross-bridges are being simulated in the context of the whole muscle, including the contribution of connective tissues, *K* should be used. For simulations at an individual myocyte level, *K*_myo_ should be used in place of *K*.

A coupled ordinary differential equation (ODE) describes each of the four state variables in Eq. 1. The rates of change of the proportion of cross-bridges in states **B** and **C** are derived from the law of mass action:

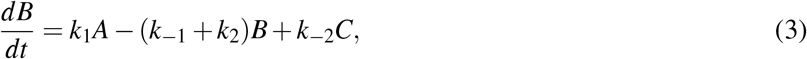

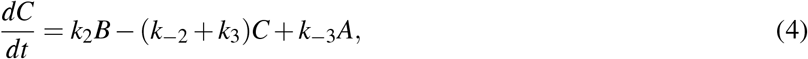

where *k*_1_, *k*_2_, *k*_3_, *k*_−1_, *k*_−2_ and *k*_−3_ are the transition rates between cross-bridge states. The proportion of cross-bridges in state **A** is given by the principle of conservation. For the maximally-activated cross-bridge model this is:

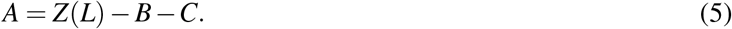

The proportion of all possible cross-bridges that are able to bind for a given sarcomere length is given by *Z* in Eq. 5. The equation for *Z* therefore describes the length-dependence in the model:

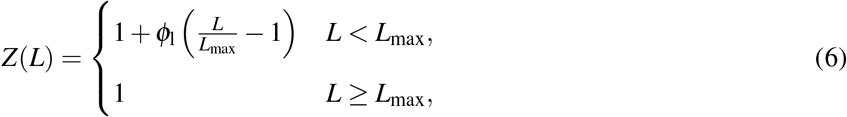

where *L* is the sarcomere length, *f*_l_ is a unitless parameter which governs the length dependence. *L*_max_ is the sarcomere length at which the maximum number of binding sites are available for cross-bridge formation and was taken as 2.3 *µ*m.

The rates of change for the mean cross-bridge strains in the attached states are governed by the following ODEs:

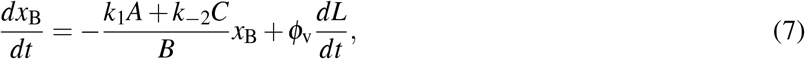

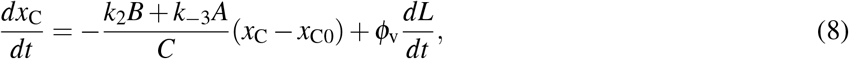

where *ϕ*_v_ is a unitless parameter governing the sarcomere velocity dependence and *x*_C0_ = 0.01 *µ*m, the strain associated with the power stroke, is the steady-state value of *x*_C_.

Most of the cross-bridge transition rates are influenced by strain or metabolite concentration. Exponential strain dependence was required on rates *k*_1_ and *k*_3_ to accurately reproduce the complex modulus. The concentration of P_i_ was included directly and scales *k*_−1_. ATP release occurs under rapid equilibrium and thus influences both *k*_−2_ and *k*_3_. These four rates are thus no longer constant parameters and are defined as follows:

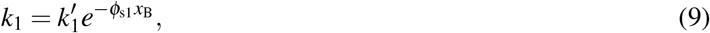

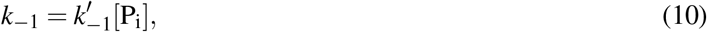

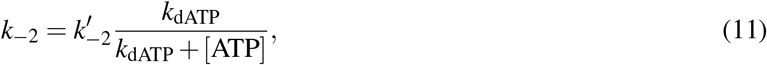

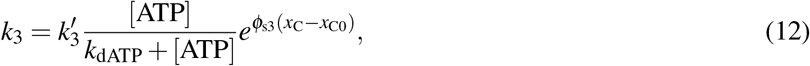

where 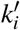 is the initial or reference value of rate *k*_*i*_, *ϕ*_s*i*_ is the parameter governing strain dependence on rate *k*_*i*_, [P_i_] is the intracellular P_i_ concentration, [ATP] is the intracellular MgATP concentration and *k*_dATP_ is the dissociation constant for MgATP from myosin.

The transition rate *k*_−3_ is often omitted from cross-bridge models for simplicity due to its negligible magnitude. We have included it here to maintain thermodynamic consistency in the cross-bridge cycle [16], and define it algebraically as:

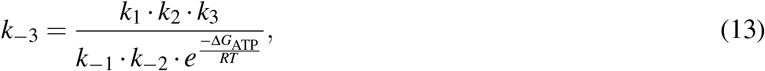

where Δ*G*_ATP_ is the free energy of ATP hydrolysis, *R* is the universal gas constant, and *T* is the temperature.

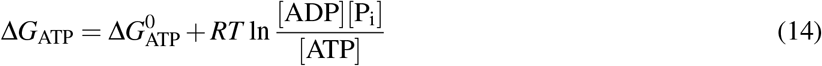

where 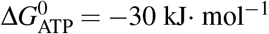 is the Gibbs free energy of ATP hydrolysis under standard conditions and [ADP] = 36 *µ*M is the intracellular ADP concentration [16].

#### 2.2.2 Model calibration

The linearised form of the model was used to identify the optimal parameters for the non-diabetic and diabetic cross-bridge models, as well as to select the optimal mechanism for strain and metabolite dependencies. The methods for linearising distortion-stiffness models are described elsewhere [14, 17]. The MATLAB particle swarm global optimiser was used to minimise the following objective function:

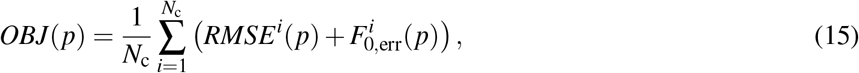

where *p* is the set of 11 variable parameters to be optimised, *N*_c_ = 5 the number of experimental conditions being simulated, *RMSE*^*i*^(*p*) is the root-mean-square error in the complex modulus prediction under parameter set *p* and condition *i* and 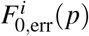 is the error in the steady-state force production under parameter set *p* and condition *i*. The *RMSE*^*i*^(*p*) is given by:

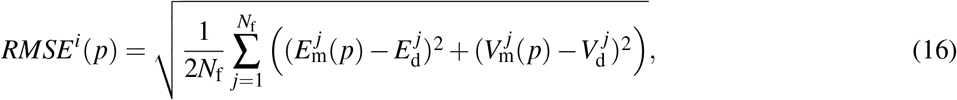

where *N*_f_ = 11, the number of frequencies at which the complex modulus was collected, *E*_m_(*p*) ^*j*^ and 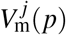 are the elastic and viscous moduli predicted by the model under parameter set *p* at frequency *j*, 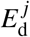 and 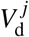 are the elastic and viscous moduli measurements at frequency *j*. The *F*^*i*^ (*p*) is given by:

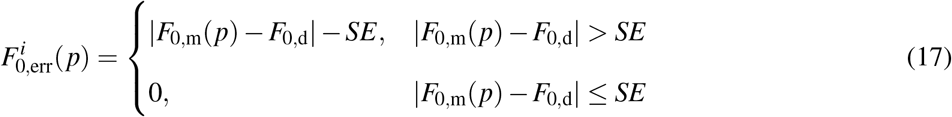

where *F*_0,m_(*p*) is the steady-state stress predicted by the model under parameter set *p, F*_0,d_ is the mean active steady-state stress from the data set and *SE* is the standard error in the specific value of *F*_0,d_. This function ensures that reasonable steady-state forces are described by the model but does not influence the objective function if the steady-state force falls within the standard error in the mean value measured.

The final non-diabetic model was fitted first and its parameters were used as a starting point for fitting the diabetic model. *K* was constrained to be less than 10000 GPa·m^−1^ as the diabetic data required a low value and too much variation between the groups in this value would result in drastically different steady-state values in the proportion of bound cross-bridges. To improve model parsimony without compromising the fit, the scaling parameter *ϕ*_x_ used in previous work [14, 18] was removed. This simplification aligned the cross-bridge strain ODEs more closely with those from the original work of Razumova et al. [15] and allowed direct measurement of the effect of the cross-bridge cycling rates on the complex modulus frequencies.

### 2.3 Muscle Model

In order to investigate the effect of diabetes on cardiac mechanoenergetics, the non-diabetic and diabetic cross-bridge models were integrated with a model of passive force and a model of thin filament activation by Ca^2+^, to create a full muscle model.

#### 2.3.1 Model of passive force

The active cross-bridge model was combined with a passive model following the approach described by Tewari et al. [19]. As depicted in Fig. 1B, this consists of a viscoelastic passive force (*F*_p1_) and a purely elastic force (*F*_p2_) in parallel with the active force generated by the cross-bridge model (*F*_a_). The total force produced by the muscle model is thus the sum of these three forces:

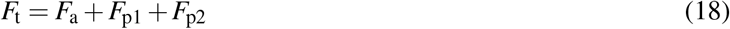

The force from the viscoelastic element is described by the following ODE:

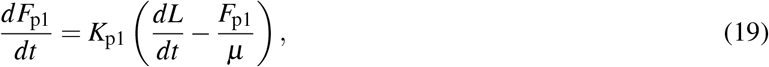

where *K*_p1_ is the constant of the series spring and *µ* is the viscosity of the dashpot. The purely elastic element was adapted from Tewari et al. [19] to reflect an exponential passive force-length relationship and was defined only above resting muscle length. It describes the steady-state passive force of the muscle as a function of length:

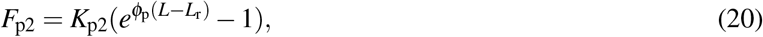

where *K*_p2_ is a scaling factor, *ϕ*_p_ is the parameter determining the curvature of the exponential relationship and *L*_r_ is the resting length of the muscle, defined in the experimental data to be 1.87 *µ*m (85 % of 2.2 *µ*m).

The four parameters of the passive force model, *K*_p1_, *µ, K*_p2_ and *ϕ*_p_, were identified by fitting to the passive complex and passive force-length relationships measured in the human atrial tissues [13]. These describe the frequency-dependent behaviour of the muscle at optimal length (Fig. 2A & B) and the length-dependent steady-state behaviour (Fig. 2C). While the complex modulus captures both elements of the passive model, the force-length relationship is required to uniquely identify the two parameters of the non-linear spring component (Eq. 20).

**Figure 2:**
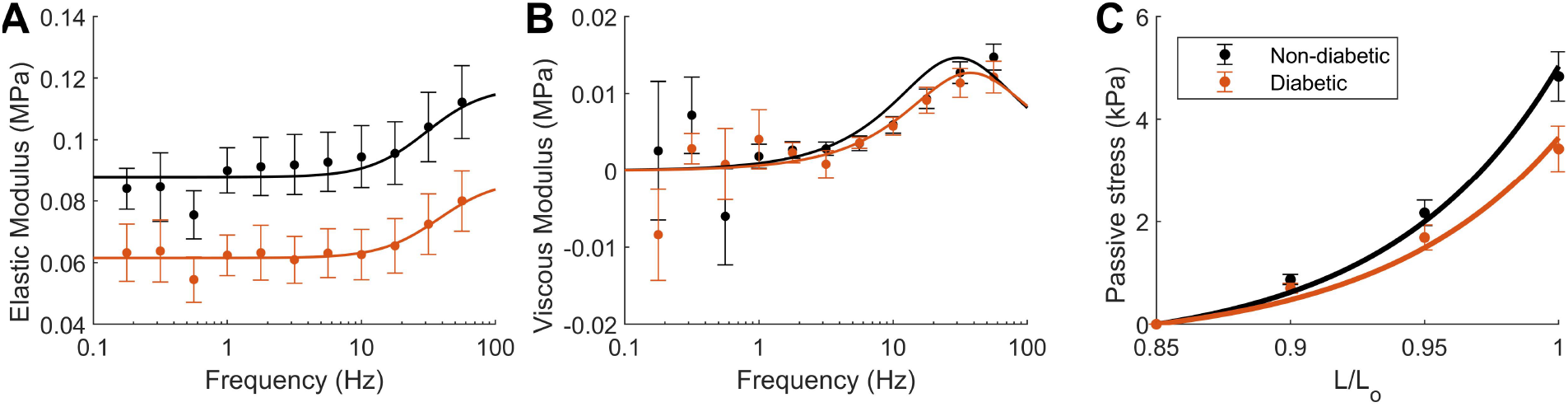
Parameterising the passive model using experimental data from human atrial tissues. Best fit of the model to the non-diabetic and diabetic passive elastic (A) and viscous modulus (B). (C) Best fit of the purely elastic force component (*F*_p2_; Eq. 20) to the non-diabetic and diabetic passive stress-length data at steady state. Filled circles with error bars represent the experimental data, and solid lines represent the fitted model.

The optimal passive model parameters are presented in Table 1. The parameter values governing the properties of the passive model are larger for the non-diabetic model, reflecting the higher passive stresses and stiffnesses measured in these trabeculae.

**Table 1:**
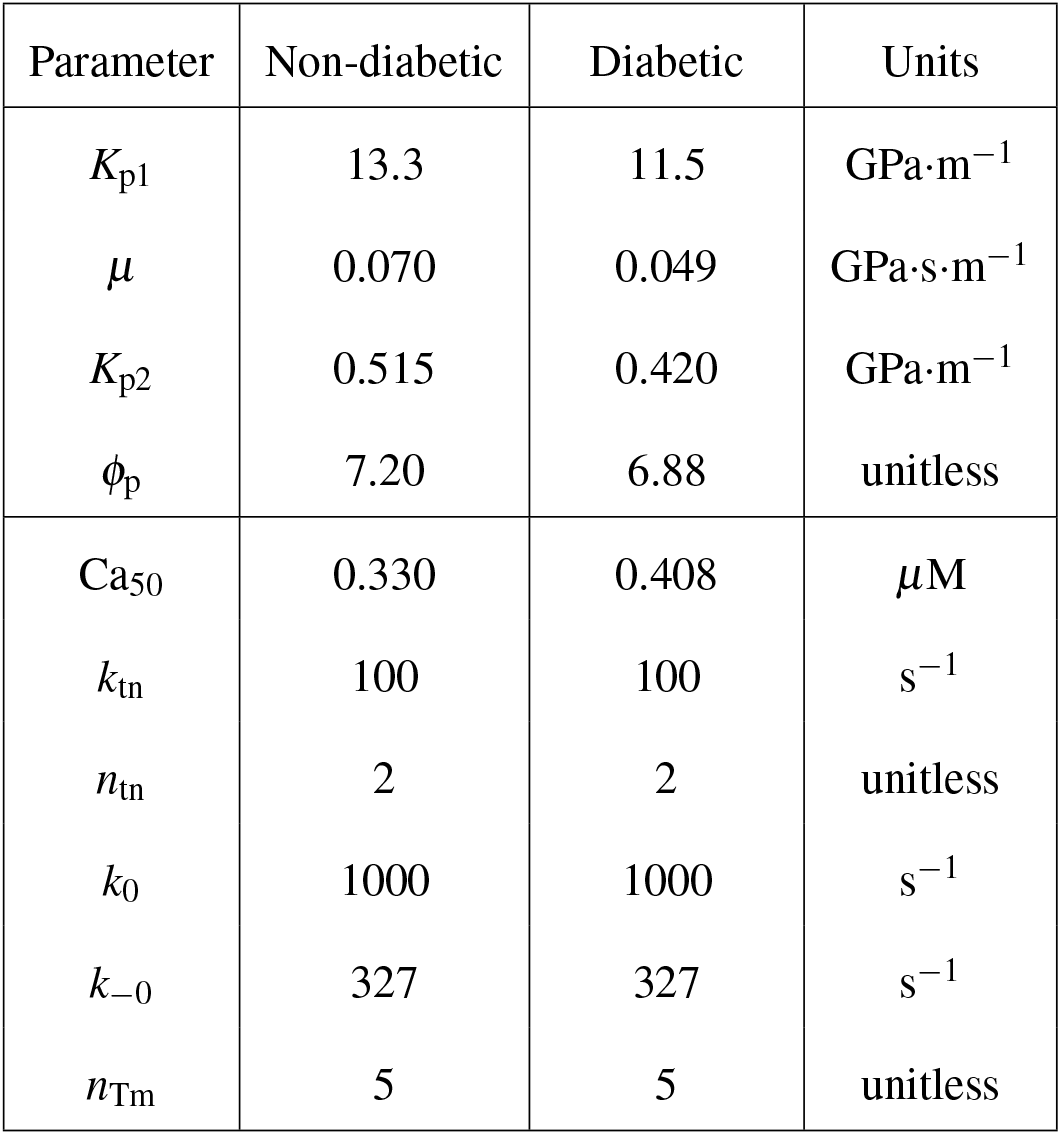
Parameters for the non-diabetic and diabetic muscle models. Upper passive model values are from fitting to the passive muscle data. Lower thin filament model values are from Land et al. [20], except for Ca_50_, which is from Jones et al. [9].

#### 2.3.2 Thin filament Ca^2+^ activation

The muscle model simulations also required a model that was sensitive to changes in intracellular Ca^2+^ concentration. We used the elements of thin filament kinetics from the Land et al. [20] human muscle model for this step, as this is a simple model which is well-suited to integration with the stiffness-distortion cross-bridge model presented in the previous section.

Fig. 1 illustrates the addition of a non-permissive state, **N**, to our cross-bridge model. The transition rates between this state and the detached state, **A**, are influenced by the proportion of regulatory units in the Ca^2+^-bound state, **CaTn**. The following equation captures the cooperative binding of Ca^2+^ to troponin:

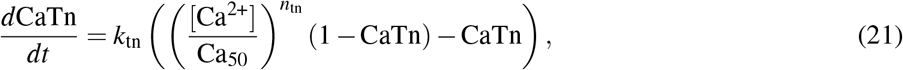

where CaTn is the proportion of TnC units with Ca^2+^ bound to their regulatory binding site, *k*_tn_ is the net unbinding rate, *n*_tn_ captures the cooperativity of the Ca^2+^-TnC binding rate, and Ca_50_ is the calcium concentration that elicits 50% activation which is commonly used to quantify the sensitivity of troponin to Ca^2+^.

Cross-bridges can transition from the non-permissive **N** state to the permissive **A** state only once the binding of Ca^2+^ moves the tropomyosin molecule and makes an actin binding site available. This process is described by the following equation:

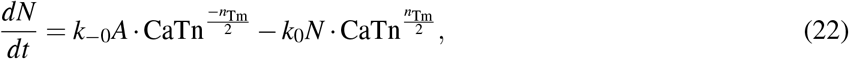

where *N* is the proportion of cross-bridges without an available actin site (in the non-permissive state), *k*_0_ and *k*_−0_ are the transition rates between these states, and *n*_Tm_ describes the cooperativity of the tropomyosin activation.

The equation describing the proportion of cross-bridges in state **A**, the detached but permissive state, was altered to include the proportion of non-permissive cross-bridges. The conservation equation previously given by Eq. 5 was modified to include State *N*:

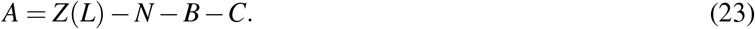

Table 1 displays the parameters used for the thin filament Ca^2+^ activation component of the muscle model. In an atrial version of the Land model [21], Ca_50_ was the only Ca^2+^ handling parameter which was altered from the original ventricular model due to the strong dependence of this parameter on the Ca^2+^ transient. In addition, there is experimental evidence of a difference in Ca^2+^ sensitivity between non-diabetic and diabetic atrial muscles [9]. Thus, we modified the values of Ca_50_ for both the non-diabetic and diabetic models based on sensitivity measurements from Jones et al. [9]. The Ca^2+^ recordings used to estimate these sensitivity values were collected from non-diabetic and diabetic human atrial trabeculae in a cohort of patients similar to those of Musgrave et al. [13]. This study was also used to inform the Ca^2+^ transients driving the muscle model simulations, which are described in further detail below.

For simplicity, the length dependence on Ca_50_ was not retained in our model, due to difficulties in direct length equivalences between the two models. As length-dependence on Ca^2+^ sensitivity is not known to be affected in diabetes, we don’t expect this to have a material influence on the comparative outputs of these model simulations.

The final step for integrating thin filament kinetics into the cross-bridge model was an adjustment to active force production in the model. The cross-bridge model was developed under the assumption that all cross-bridges are in the permissive state at pCa 4.5 ([Ca^2+^] = 32 *µ*M). While troponin is saturated at this level of Ca^2+^, the description of transitions between **N** and **A** means that there will still be a small, constant fraction (5 % to 10 % for the rate constants in this model) of cross-bridges that remain in the non-permissive state. This causes a reduction in the maximum force produced by the cross-bridge at maximal Ca^2+^ concentrations, as not all possible cross-bridges are available for recruitment. To account for this, we adjusted the value of *K* for each model such that the muscle model produced the original maximal force under an input value of [Ca^2+^] = 32 *µ*M. This adjustment can be thought of as rescaling the total number of cross-bridges in the muscle.

### 2.4 Muscle simulations

#### 2.4.1 General simulation methods

Simulations of non-diabetic and diabetic muscle models were performed in MATLAB using the ode15s solver. Unless stated otherwise, the parameters listed in Tables 1 and 3 and the baseline values of ATP and P_i_ (5 mM and 1 mM) were used for the simulations.

The muscle models were driven by Ca^2+^ transients based on measurements from non-diabetic and diabetic human atrial trabeculae contracting at 1 Hz at body temperature [9]. For the non-diabetic Ca^2+^ transient, we scaled a generic human atrial Ca^2+^ transient to match the amplitude and diastolic values of those reported for non-diabetic trabeculae by Jones et al. [9]. For the diabetic Ca^2+^ transient, the amplitude, diastolic level, time to peak and decay constant were scaled by the proportion that each of these values were measured to differ from the non-diabetic group. The two transients used in the simulations are shown in Fig. 3. As found in Jones et al. [9], the diabetic transient has a prolonged rise time and relaxation rate, with an elevated diastolic Ca^2+^ concentration.

**Figure 3:**
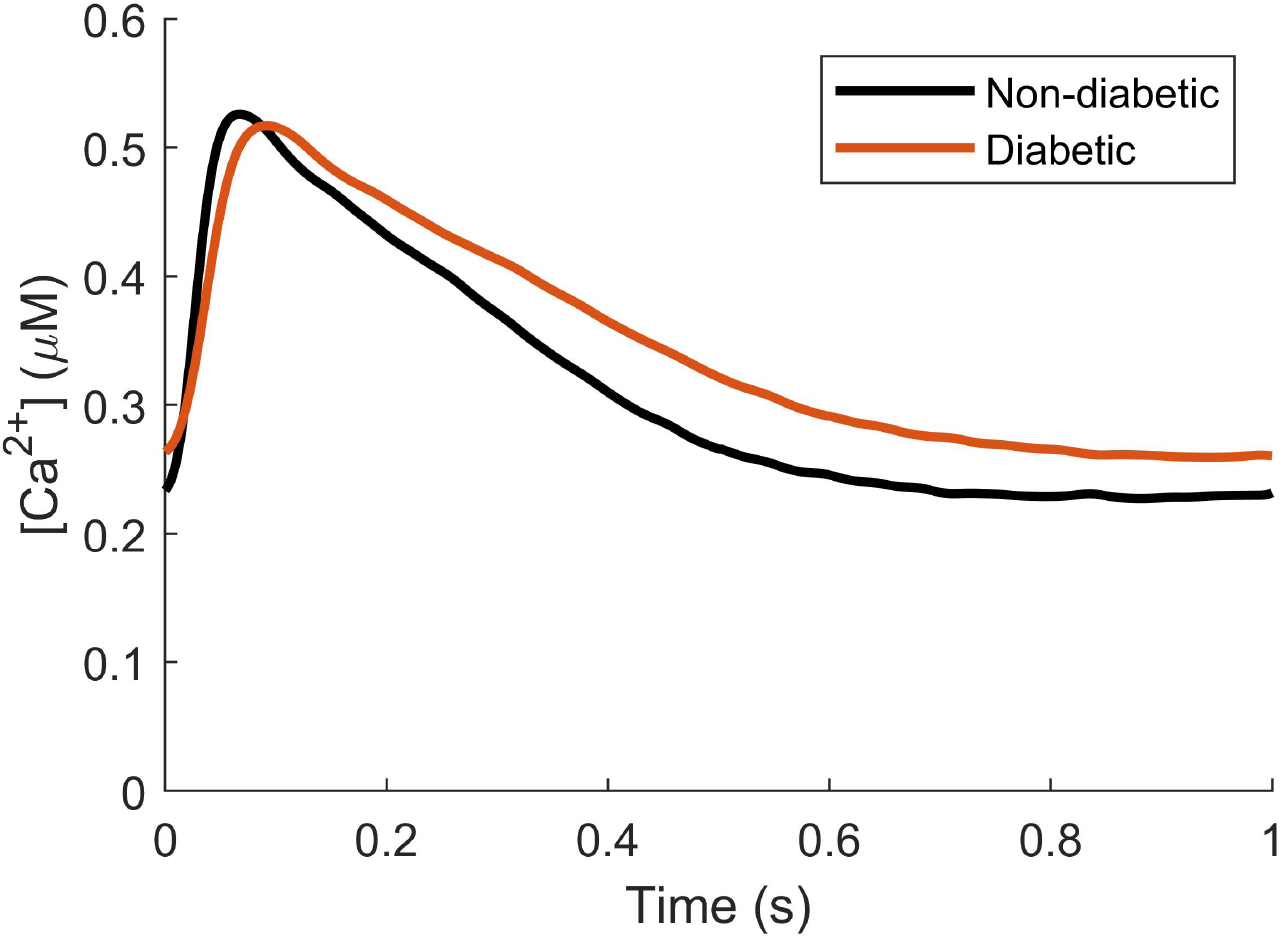
Ca^2+^ transients used to drive the muscle models. Non-diabetic and diabetic Ca^2+^ transients were derived from human atrial trabeculae contracting at 1 Hz at body temperature [9]. The Ca^2+^ sensitivity values, Ca_50_, for the non-diabetic and diabetic groups are 0.330 *µ*M and 0.408 *µ*M, respectively.

#### 2.4.2 Isometric contraction

In isometric contraction simulations, the muscle length was held fixed at a sarcomere length of 2.2 *µ*m and activated by a 1 Hz Ca^2+^ transient (Fig. 3). The simulation was run until a beat-to-beat steady state was reached. The diastolic (minimum) and systolic (maximum) stresses were found, as well as the twitch amplitude. The twitch duration was calculated as the time it took for the stress to return to 5 % of the twitch amplitude after initially rising above this value.

#### 2.4.3 Work-loop contraction

Work-loop contraction simulations were performed to predict how diabetes-driven differences in Ca^2+^ transients and cross-bridge function affected muscle mechanoenergetics under loading conditions similar to those experienced by the heart *in vivo*. Stress-length work-loops are a one-dimensional analogue to the pressure-volume loop of the three-dimensional heart. Although work-loop contractions are characteristic of tissues in the ventricular walls, using an atrial model in this context allows investigation of the general effect of diabetes on cardiac tissue mechanoenergetics. We assume that diabetes affects atrial and ventricular tissues similarly; as such our findings are also applicable to ventricular tissues. These work-loop simulation protocols mimic the experimental protocols that are used for cardiac trabeculae [22, 23].

The simulation is driven by a 1 Hz Ca^2+^ transient and starts at an initial sarcomere length of 2.2 *µ*m. The first phase of the simulation is isometric tension development, as rising Ca^2+^ concentration drives the generation of active stress. Once the muscle stress reaches the prescribed afterload, an isotonic phase begins where the muscle shortens against the constant stress. Isometric relaxation occurs when the muscle can no longer continue to shorten and the muscle is finally restretched back to the intial sarcomere length before the end of the 1 Hz cycle. These simulations were repeated at 10 prescribed afterloads which range between the diastolic stress and the systolic stress produced by the isometric twitch. As with the isometric contractions, these work-loop simulations were run until a beat-to-beat steady state was reached.

Several metrics of contraction were found from the series of work-loops performed with each muscle model. The work done throughout a given work-loop was found by numerically integrating the stress with respect to sarcomere length, reflecting the area within the stress-length loop. The extent of shortening reflected the end-systolic length for each afterload. The maximum velocity of shortening was found by taking the minimum (largest negative) gradient of the sarcomere length output and normalised to muscle length. Maximum velocity of shortening also allowed for the calculation of maximum power of shortening, the product of the maximum velocity of shortening and the given afterload.

In addition, we performed an analysis to quantify the energy utilised by the muscle throughout the contraction. These methods have been adapted from Tran et al. [24]. Each cross-bridge is assumed to consume 1 ATP. The rate of ATP consumption, *J*_xb_ in this model is given by:

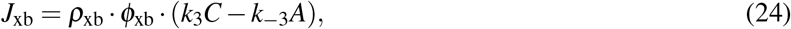

where *ρ*_xb_ is the density of cross-bridges, *ϕ*_xb_ is the number of ATP molecules consumed per cross-bridge cycle (assumed to be 1), and the final term describes the net transition of cross-bridges from state **C** to state **A**. We multiplied the value of *ρ*_xb_ = 0.25 mol·m^−3^ used by Tran et al. [24] by the myocyte density in each group (*ρ*_myo_) to account for the measured density of myofilaments in the human trabeculae [13].

The total amount of ATP energy consumed during a single work-loop is given by the time integral of this consumption rate multiplied by the Gibbs free energy of ATP:

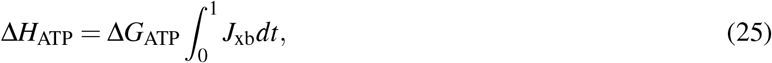

where Δ*H*_ATP_ is the total change in enthalpy per loop and Δ*G*_ATP_ is calculated using Eq. 14, assuming [ADP] = 0.036 mM.

The change in enthalpy per twitch was also used to quantify the cross-bridge efficiency, by dividing the work done in a given work-loop by the ATP energy consumed.

#### 2.4.4 Varying Pi and ATP

Isometric contraction simulations were performed under high (10 mM) P_i_ and low (1 mM) ATP to quantify the effects of these metabolites on the morphology of the stress twitch time course. Work-loop contraction simulations were also performed under a range of P_i_ and ATP concentrations to characterise the effects of these metabolites on mechanoenergetic metrics described in Section 2.4.3. Raised P_i_ and reduced ATP conditions were selected as these are the trends in metabolite availability seen in diabetic hearts [6, 7].

#### 2.4.5 Cross-over simulations

To investigate the relative effects of diabetes-induced alterations on Ca^2+^ handling compared to those on cross-bridge function, four distinct sets of simulations were performed. In the first two sets, non-diabetic and diabetic Ca^2+^ transients were matched to their respective non-diabetic and diabetic cross-bridge models. In the remaining two sets of cross-over simulations, diabetic Ca^2+^ transients were paired with the non-diabetic cross-bridge model, and vice versa. The value of Ca_50_ was always matched to the Ca^2+^ transient used for the Ca^2+^ handling. The four sets of simulations were all performed with the same passive model (non-diabetic) in order to control for differences in passive stress and isolate the effects of diabetes on muscle activation and contraction.

#### 2.4.6 Parameter sensitivity analysis

Finally, to systematically examine the effect of the parameters within each of the muscle model components on the outputs of the simulations, we performed a sensitivity analysis on all of the model parameters. A range of output metrics from the muscle model simulation outputs were computed under baseline metabolite concentrations using the non-diabetic model and compared to simulations where each parameter was increased by 10 %. The altered parameters were the 11 fitted cross-bridge model parameters (Table 3), the four passive model parameters and the six thin filament parameters (Table 1). To consider the effect of changing the input Ca^2+^ transient, we also increased the diastolic Ca^2+^, the systolic Ca^2+^ and the duration of the twitch by 10 %. The simulation outputs compared were diastolic stress, systolic stress, twitch amplitude and twitch duration for the isometric twitch and the maximum value for each of the outputs from the work-loop simulations (i.e. maximum work, shortening extent, shortening velocity, shortening power, enthalpy and cross-bridge efficiency).

### 2.5 Model availability

Complete model code with all relevant parameters is available at https://github.com/JuliaMusgrave/AtrialModel_2025_Human. Complete muscle model equations and parameters are listed for convenience in the Section 4.5.

## 3 Results

### 3.1 Cross-bridge model

The optimal cross-bridge model fits for both the non-diabetic and diabetic datasets produced steady-state stresses within the bounds of the standard errors under all five metabolite conditions. The model predictions of these stresses are presented for comparison with experimental data from Musgrave et al. [13] in Table 2.

**Table 2:** Simulation of steady-state active stress using the non-diabetic and diabetic models for different metabolite conditions. Simulation values are compared to the experimental data that were used in the parameter optimisation process.

|  | Non-diabetic stress (kPa) |  | Diabetic stress (kPa) |  |
| --- | --- | --- | --- | --- |
|  | Exp Data | Model | Exp Data | Model |
| Baseline | $20.4 \pm 1.8$ | 22.0 | $16.2 \pm 2.1$ | 16.9 |
| 1 mM ATP | $21.5 \pm 2.7$ | 23.1 | $19.0 \pm 3.1$ | 18.4 |
| 0.1 mM ATP | $24.9 \pm 2.8$ | 27.2 | $18.9 \pm 2.3$ | 19.1 |
| 0 mM $P_i$ | $20.9 \pm 2.4$ | 22.6 | $17.1 \pm 2.2$ | 17.2 |
| 10 mM $P_i$ | $17.0 \pm 1.6$ | 17.9 | $12.6 \pm 1.8$ | 14.4 |

**Table 3:** Parameter values for the non-diabetic and diabetic cross-bridge models found by fitting to active muscle data. Values for *ρ*_myo_ were taken directly from Musgrave et al. [13] and subsequently used to calculate *K*_myo_ according to Eq. 2. The other 11 parameters were found from the calibration protocol described in Section 2.2.2.

| Parameter | Non-diabetic | Diabetic | Units |
| --- | --- | --- | --- |
| $k'_1$ | 89.0 | 44.3 | $s^{-1}$ |
| $k'_{-1}$ | 6.68 | 2.44 | $s^{-1}$ |
| $k_2$ | 88.8 | 86.27 | $s^{-1}$ |
| $k'_{-2}$ | 113 | 93.57 | $s^{-1}$ |
| $k'_3$ | 96.6 | 50.00 | $s^{-1}$ |
| $\phi_v$ | 0.029 | 0.036 | unitless |
| $\phi_l$ | 5.00 | 5.00 | unitless |
| $K$ | 9030 | 5270 | $\text{GPa}\cdot\text{m}^{-1}$ |
| $\rho_{\text{myo}}^*$ | 0.361 | 0.292 | unitless |
| $K_{\text{myo}}^*$ | 25000 | 18000 | $\text{GPa}\cdot\text{m}^{-1}$ |
| $\phi_{s1}$ | 257 | 7.13 | $\mu\text{m}^{-1}$ |
| $\phi_{s3}$ | 44.9 | 199 | $\mu\text{m}^{-1}$ |
| $k_{\text{dATP}}$ | 0.281 | 5.00 | mM |

As all of the model predictions were within the standard error of the steady-state active stress measurements, the value of the optimal objective function (Eq. 15) simply reflected the RMSE of the complex moduli. The best-fit model to the non-diabetic data provided an objective function value of 12.5 kPa or a normalised RMSE of 2.99 %. The best-fit model to the diabetic data provided an objective function value of 10.6 kPa or a normalised RMSE of 3.24 %. These models and their fit to the experimental data (complex moduli) are shown in Fig. 4.

**Figure 4:**
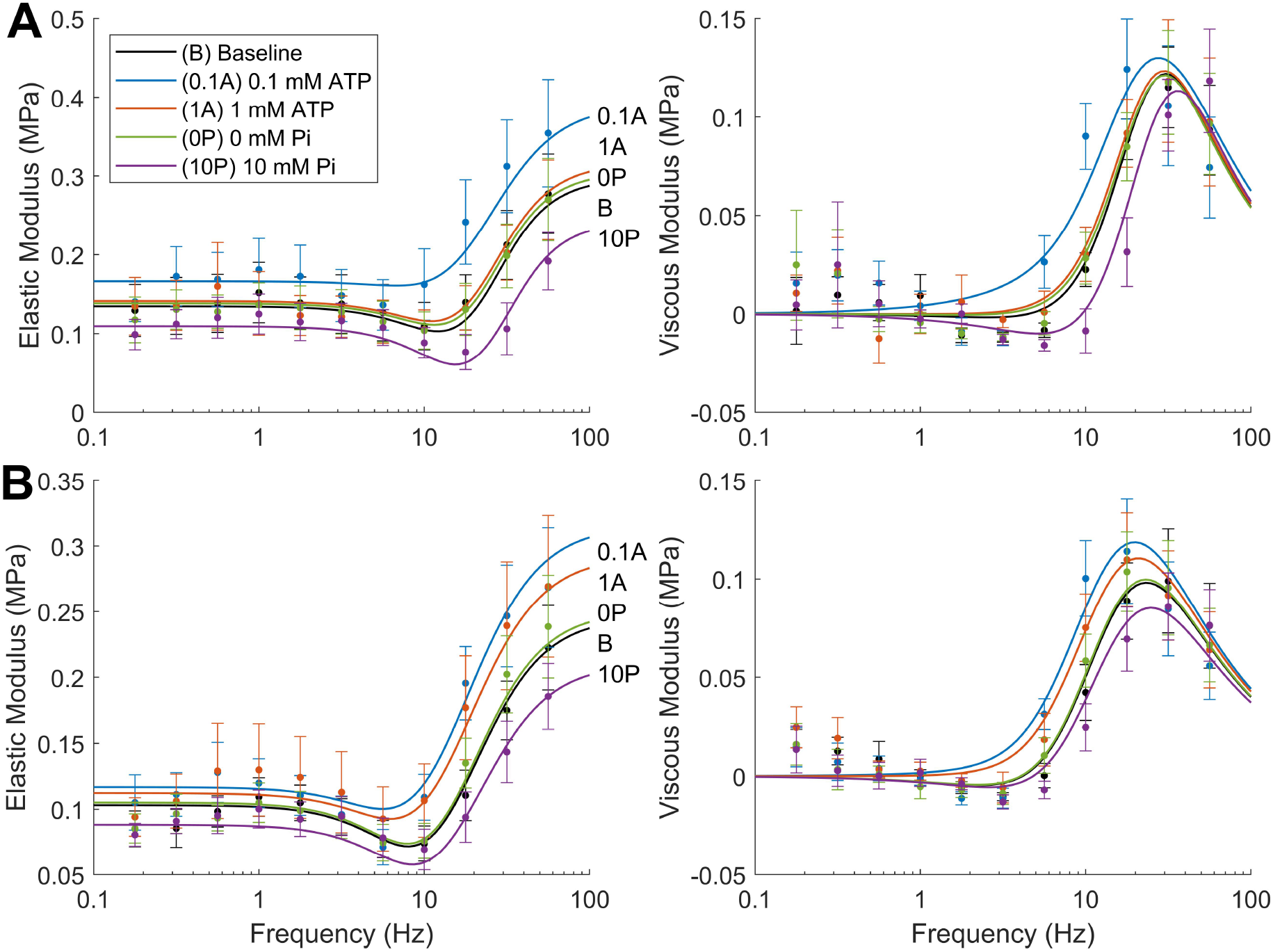
Parameterising the cross-bridge models. Best fit of the cross-bridge models to non-diabetic (A) and diabetic (B) human active complex modulus data from Musgrave et al. [13] under different metabolite concentrations.

To obtain these two model fits of similar goodness for significantly different sets of data, there were inevitable differences in the cross-bridge parameters between the non-diabetic and diabetic models. The parameters for the respective models are presented in Table 3. The variation in optimal cross-bridge parameters for the non-diabetic and diabetic models demonstrate clearly the mechanisms that underlie the diabetes-induced differences seen in the experimental measurements. The optimal fit resulted in higher values for the cross-bridge transition rate parameters in the non-diabetic model compared to the diabetic model. However, significantly different values of *k*_dATP_ mean that *k*′_−2_ and *k*_3_ do not reflect the values of their respective transition rates in the model (see Eqs. 11 and 12). When these are evaluated at the baseline ATP concentration (5 mM), *k*_−2_ is faster in the diabetic model (46.8 s^−1^ vs 6.0 s^−1^) while *k*_3_, along with the other rates, is slower (25.0 s^−1^ vs 91.4 s^−1^). Since the detachment rates (*k*_3_ and *k*_−1_) in particular are slower in the diabetic model, cross-bridges are more likely to be attached, resulting in the diabetic model having a higher value of *C*_0_ (the steady-state proportion of cross-bridges bound in the post-power stroke state) at baseline levels (0.32 vs 0.24). Relating to this, *K* is much lower in the diabetic model, which allows lower steady-state stresses to still be produced despite a larger proportion of attached cross-bridges. Decomposing this value according to Eq. 2 shows that this reflects reduced stiffness at the myocyte level (*K*_myo_) as well as reduced myocyte density (*ρ*_myo_) measured in the diabetic muscles [13].

As the P_i_ concentration was incorporated directly into *k*_−1_, the higher value of *k*′_−1_ in the non-diabetic model reflects a higher sensitivity to P_i_. On the other hand, since the rapid equilibrium formulation was used for ATP binding, its sensitivity is primarily modulated through the value of *k*_dATP_. The higher value of this parameter in the diabetic model reflects an increased sensitivity of the model to ATP within this physiological range.

### 3.2 Isometric Contraction

When the non-diabetic and diabetic muscle models were activated by their respective Ca^2+^ transients, there were a number of apparent differences in the simulated isometric twitches (Fig. 5A). The diastolic stress was higher in the non-diabetic muscles, reflecting the increased passive stresses measured in these trabeculae and consequent higher passive stiffness parameters in this model (Table 1). The systolic stress and twitch amplitude were 41 % and 45 % lower for the diabetic model, respectively (20.2 kPa vs 12.0 kPa for the systolic stresses and 14.5 kPa vs 7.9 kPa for the twitch amplitudes). Finally, the twitch duration was 23 % longer for the diabetic model (674 ms vs 550 ms duration at 95 % relaxation from the peak).

**Figure 5:**
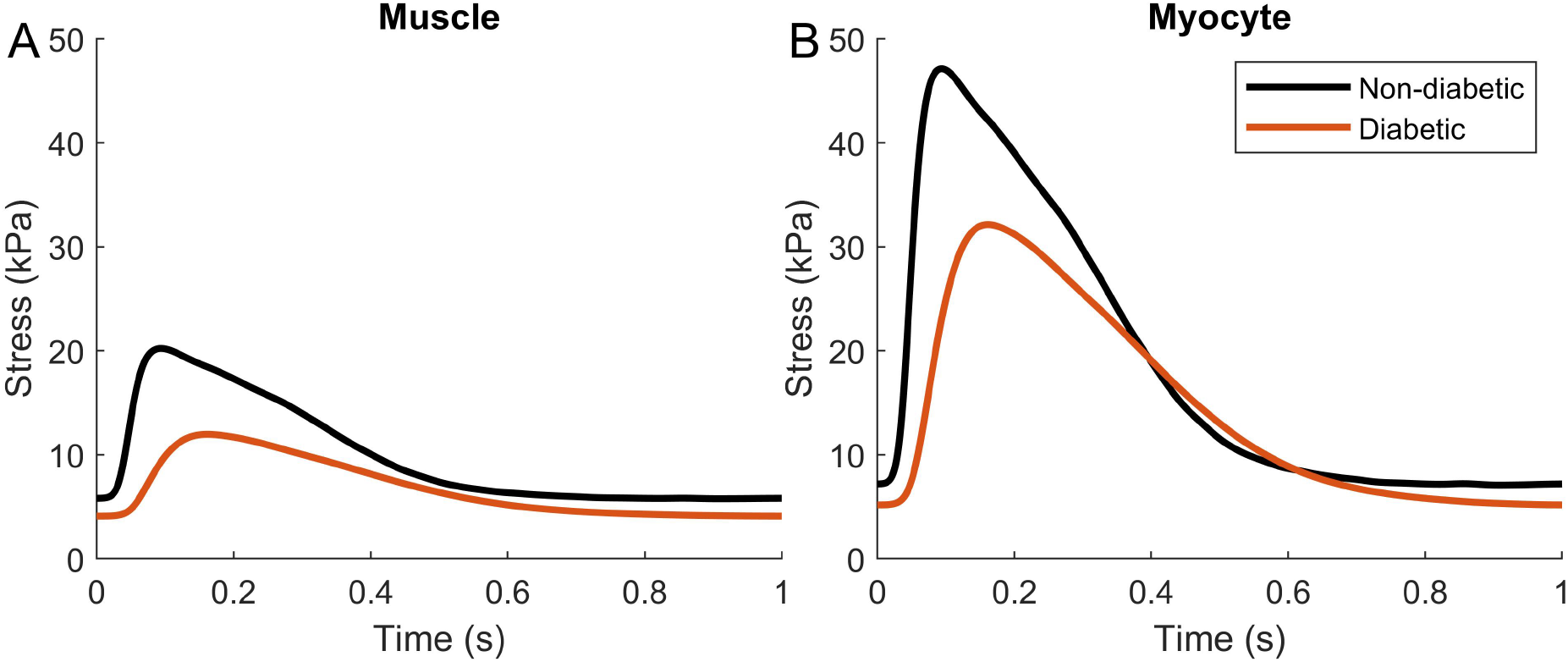
Model simulation of isometric contractions. Isometric twitches were simulated using the non-diabetic and diabetic muscle models activated by their respective Ca^2+^ transients and Ca^2+^ sensitivity values. The twitch force is normalised to either the trabeculae cross-sectional area to produce muscle stress (A) or to the myofilament area to produce myocyte stress (B).

For comparison to the isometric twitches that might be seen within an individual myocyte, we repeated the simulation using *K*_myo_ (Eq. 2) instead of *K* to scale the force produced by the cross-bridges (Fig. 5B). As the trabeculae are approximately one-third myofilament [13], this resulted in much larger twitch amplitudes compared to the muscle model (amplitudes of 40.0 kPa and 26.9 kPa for the non-diabetic and diabetic models, respectively). Due to their differences in myofilament fraction, the amplitudes in the non-diabetic and diabetic models are more similar to each other at the myocyte level than at the muscle level (33 % less in the diabetic myocyte model, as opposed to 45 % in the muscle model). The twitch duration does not change with the scaling in the myocyte model, so the difference appears more pronounced than in the muscle model.

### 3.3 Work-loop contraction

The time courses for muscle stress and length during work-loop contraction simulations at different loads are shown in Fig. 6. The isometric twitches, arising from an afterload against which the muscle cannot shorten, are illustrated in bold. The relative afterloads depicted on the right-side y-axes were normalised to the peak stress of the isometric contraction. A parametric plot of the stress and length time courses produces the work-loops in the bottom panels.

**Figure 6:**
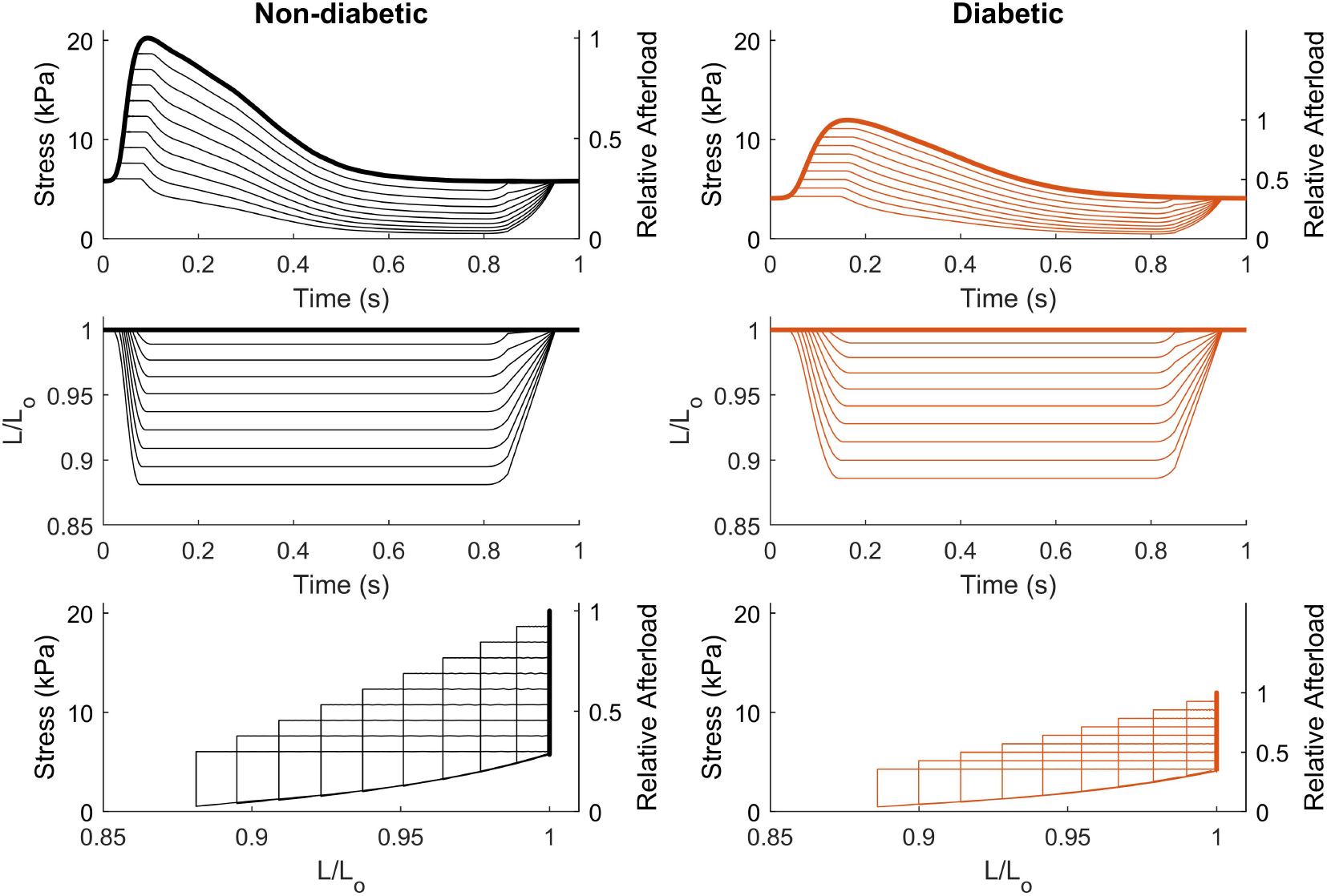
Model simulations of work-loop contractions. Work-loops were simulated for the non-diabetic and diabetic muscle models over a range of afterloads (expressed relative to the peak isometric stress). The bold lines indicate the work-loop performed at the maximum afterload (a relative afterload of 1), which is the isometric contraction. The minimum afterload is in the vicinity of passive stress, i.e., zero active stress. Stress and length (relative to the optimal length, L_0_) time courses are plotted in the upper four panels. The lower panels show stress and length plotted parametrically to reveal the work-loops generated by these contractions.

When analysing the output metrics of the work-loop simulations as a function of relative afterload, differences in the mechanoenergetics of these muscles were revealed (Fig. 7). The maximum work generated during work loop contraction simulations (area within the work-loops shown in lower panel of Fig. 6) was 44 % lower for the diabetic muscles (Fig. 7A). The extent of shortening was similar across afterloads for both muscle models (Fig. 7B), so the difference in work was entirely due to the difference in force generation. The maximum shortening velocity was 52 % lower in the diabetic muscle model (Fig. 7D). Combined with lower absolute afterload stresses, this resulted in maximum shortening power generation by the diabetic model being diminished by 69 % (Fig. 7E).

**Figure 7:**
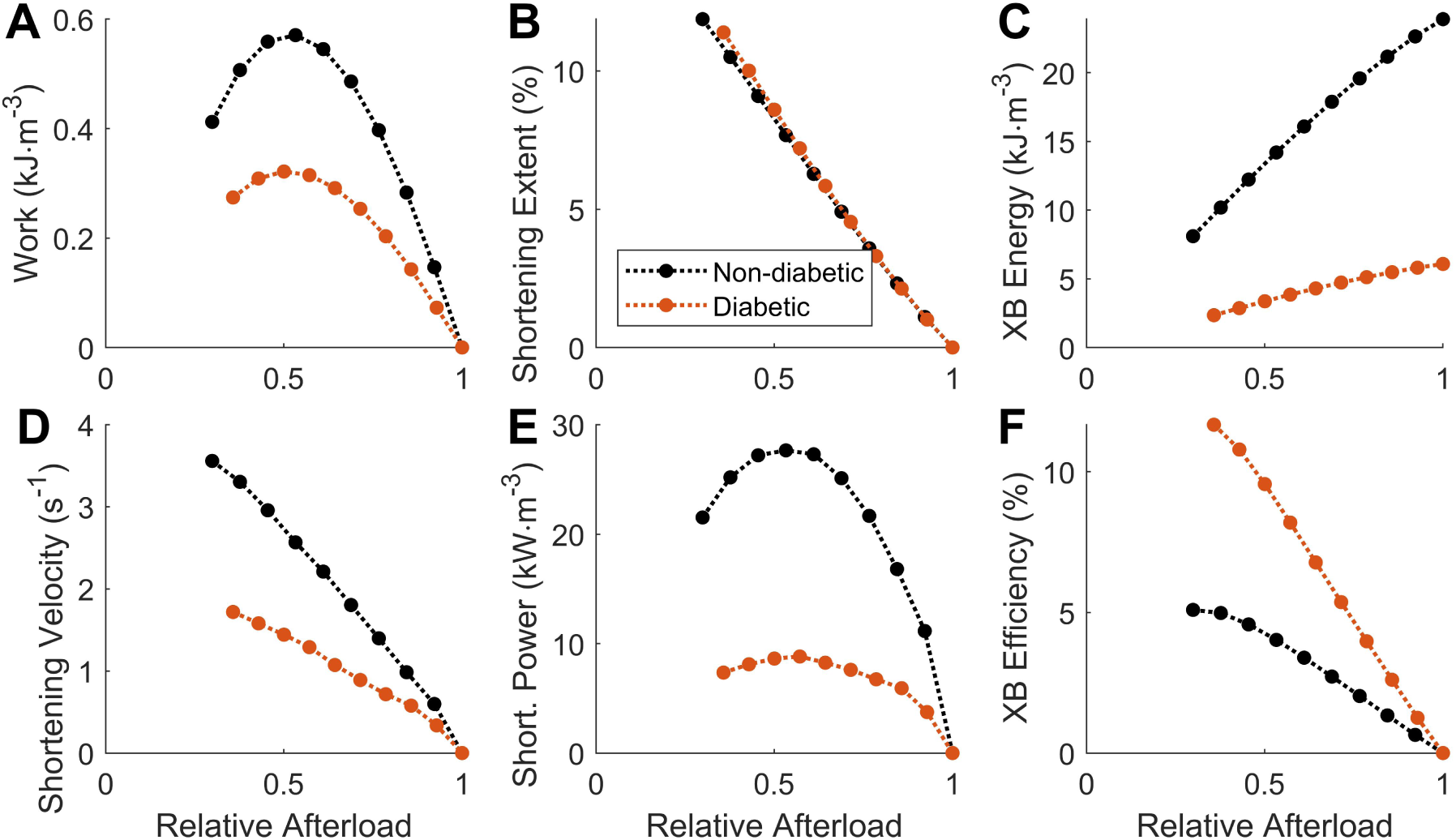
Work-loop mechanoenergetic outputs from the non-diabetic and diabetic muscle model simulations as a function of relative afterload. (A) Work done by the muscle during each work-loop contraction. (B) Percentage of muscle length shortened during each work-loop. (C) Energy consumed by cross-bridges (XB), calculated by the change in XB enthalpy per twitch. (D) Maximum velocity of shortening during work-loop contractions. (E) Maximum power of shortening, calculated as the product of shortening velocity and stress. (F) Efficiency of work done by the cross-bridges, calculated as the ratio of work to change in XB enthalpy during a twitch.

The peak energy consumed by cycling cross-bridges was reduced by 75 % in the diabetic model (Fig. 7C). Although the diabetic model generated less work, the decrease in energy consumption was greater, leading to a higher cross-bridge efficiency in the diabetic model. The peak cross-bridge efficiency in the diabetic model was more than two times higher compared to that of the non-diabetic model (Fig. 7F).

### 3.4 Effect of metabolite concentrations

The next set of results explores the effects of altered metabolite concentrations on isometric and work-loop simulations. The conditions of elevated P_i_ and lowered ATP concentrations were of particular interest, as they are most likely to occur in the hearts of individuals with type 2 diabetes [6, 7].

#### Raised P_i_ concentration

Isometric twitch simulations were performed at 10 mM P_i_ and compared against previous simulations where P_i_ was set at 1 mM (Fig. 8A). ATP was kept at 5 mM for these simulations. For both non-diabetic and diabetic models, the increase in P_i_ caused a reduction in the isometric twitch amplitude. This follows from the mechanistic effect of P_i_ that reduces the net attachment rate of cross-bridges. Proportionally, the decrease in magnitude of the twitch amplitude was similar between the two models (26 % vs 28 %).

**Figure 8:**
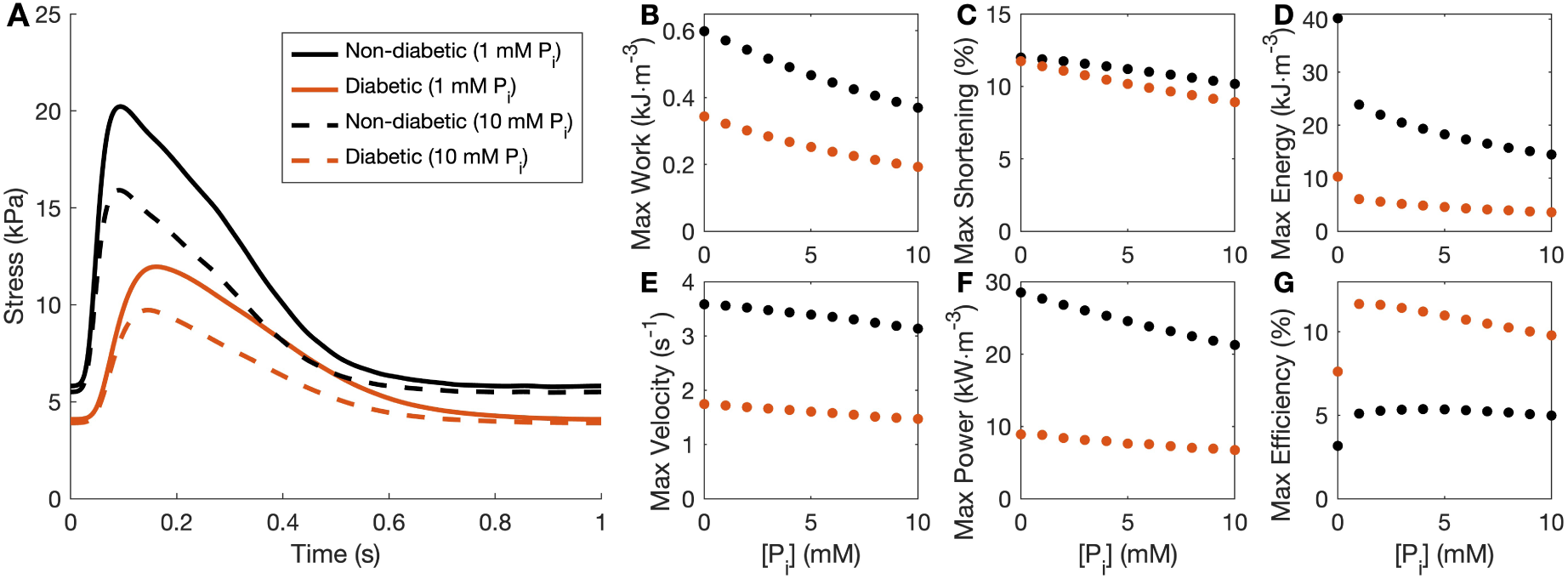
Effect of P_i_ on isometric and work-loop contractions. (A) Isometric twitches simulated using the non-diabetic (black) and diabetic (orange) muscle models with baseline (1 mM, solid lines) and high (10 mM, dashed lines) P_i_ concentrations. (B-G) Maximum output values of the six work-loop metrics plotted in Fig. 7 as a function of P_i_ concentration.

In work-loop simulations, increasing P_i_ concentration resulted in decreasing values for all of the outputs we quantified in both the non-diabetic and diabetic models (Fig. 8B-G). Higher P_i_ led to prominent reductions in work, enthalpy and shortening power. These metrics, especially power and enthalpy, were less sensitive to P_i_ in the diabetic model than in the non-diabetic model.

#### Lowered ATP concentration

Isometric simulations were performed at 1 mM ATP and compared against previous simulations where ATP was set at 5 mM (Fig. 9A). P_i_ was kept at 1 mM for these simulations. Lowering ATP resulted in a 9 % increase in the twitch amplitude in the non-diabetic model, and a 29 % increase in the diabetic model. There was also an increase in diastolic stress in the diabetic model which was not observed in the non-diabetic model. The increase in twitch duration was also greater in the diabetic model (2 % vs 25 %). Mechanistically, lower ATP reduces the rate of cross-bridge detachment [14]. This leads to an increase in the amplitude of the twitch and a decrease in the rate of relaxation, which prolongs the duration of the twitch and increases diastolic stress. The more prominent effect of a reduction of ATP concentration from 5 mM to 1 mM in the diabetic model aligns with the greater sensitivity to ATP, evidenced by the high *k*_dATP_ value obtained from model fitting to the data (Table 3). The increased sensitivity reflects the lack of saturation of ATP at these concentrations in the diabetic trabeculae [13].

**Figure 9:**
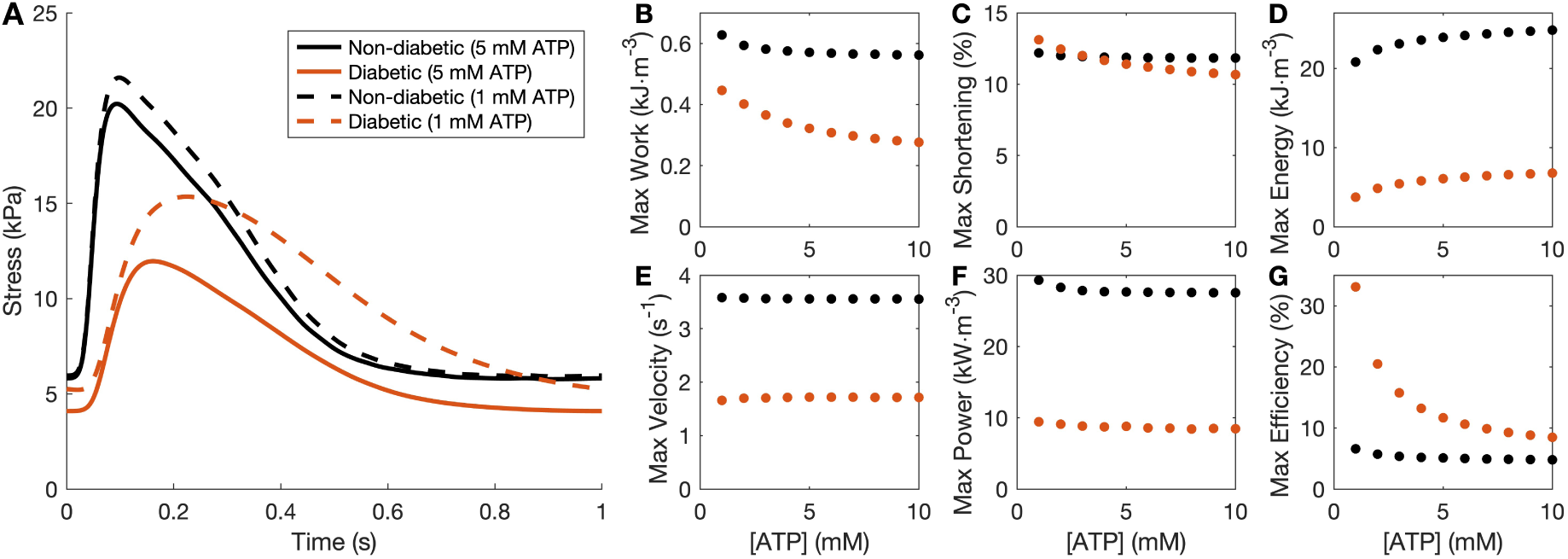
Effect of ATP on isometric and work-loop contractions. (A) Isometric twitches simulated using the non-diabetic (black) and diabetic (orange) muscle models with baseline (5 mM, solid lines) and low (1 mM, dashed lines) ATP concentrations. (B-G) Maximum output values of the six work-loop metrics plotted in Fig. 7 across ATP concentrations between 1 mM and 10 mM.

Lower ATP concentrations increased maximum work production in both simulations, with a more pronounced change in the diabetic model (Fig. 9B). There was a small increase in the maximum extent of shortening in the diabetic model, but no appreciable change in the non-diabetic model (Fig. 9C). The diabetic model also showed a small decrease in maximum shortening velocity with the lower ATP concentration, while there was no change in the non-diabetic model (Fig. 9D). On the other hand, maximum shortening power increased slightly with a more pronounced change in the non-diabetic model (Fig. 9F). The rate of ATP consumption is directly affected by its concentration, so the enthalpy per twitch increased with ATP to a similar degree for both models (Fig. 9D). As more work was being done for less energy expenditure at low ATP concentrations, this resulted in an increase in cross-bridge efficiency for both models (Fig. 9G). This was more pronounced in the diabetic model, due to its larger increase in work.

### 3.5 Model sensitivity analysis

#### 3.5.1 Cross-over simulations

To identify the relative contributions of diabetes-driven changes in Ca^2+^ handling and cross-bridge function, an additional set of two cross-over simulations was performed. These consisted of simulations where the diabetic Ca^2+^ handling was paired with the non-diabetic cross-bridge model and vice versa. Fig. 10 reproduces the non-diabetic and diabetic model simulations (Figs. 6 and 7) in black and orange solid lines, respectively, but here, the passive force model is kept the same (non-diabetic). The new cross-over simulations are depicted in green dashed and purple dotted lines. When comparing the non-diabetic (black solid) and diabetic (orange solid) models, the only qualitative effect of using the same passive force model was the elimination of a difference in diastolic stress.

**Figure 10:**
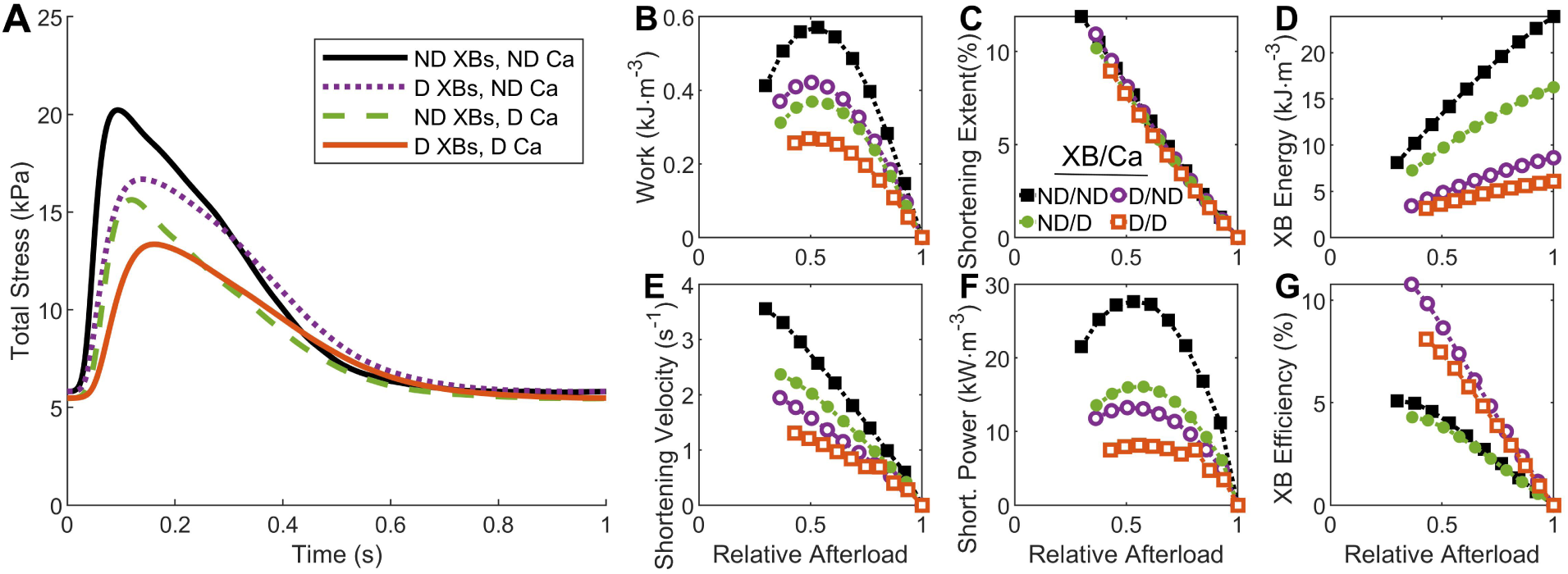
Effect of diabetes-induced changes in Ca^2+^ handling and cross-bridge function on muscle model simulations. (A) Isometric twitches simulated using the non-diabetic model (black solid line), the diabetic model (orange solid line), diabetic Ca^2+^ handling paired with the non-diabetic cross-bridge model (green dashed line) and non-diabetic Ca^2+^ handling paired with the diabetic cross-bridge model (purple dotted line). (B-G) Work loop outputs (as in Fig. 7) for the same four model configurations. Filled black squares represent the non-diabetic model simulation (ND/ND); unfilled orange squares represent the diabetic model simulation (D/D); filled green circles represent the simulation with the diabetic Ca^2+^ handling and the non-diabetic cross-bridge model (ND/D); and unfilled purple circles represent the simulation with the non-diabetic Ca^2+^ handling and the diabetic cross-bridge model (D/ND).

In isometric simulations, it is evident that diabetic Ca^2+^ handling and diabetic cross-bridges contributed almost equally to the reduced amplitude in the diabetic model (Fig. 10A). Compared to the non-diabetic model (black solid), the diabetic model (orange solid) had a twitch amplitude that was 45 % lower. When only the Ca^2+^ handling (green dashed) or the cross-bridge model (purple dotted) was diabetic, the amplitude of the twitch reduced by a similar amount, between 25 % and 30 %.

The increase in the duration of the twitch in the diabetic model was primarily due to the diabetic cross-bridges. Compared to the non-diabetic model, the duration of the twitch increased by 23 % in the diabetic model (orange solid). When only the Ca^2+^ handling was diabetic (green dashed), the increase in the duration of the twitch was only 7 % compared to an increase of 15 % when only the cross-bridge model was diabetic (purple dotted). This is also evident from the plot, qualitatively, as the two curves with diabetic cross-bridges (orange solid and purple dotted lines) both took longer to reach their peak stress and exhibited a less angular morphology.

For many of the work-loop outputs, the differences observed between the diabetic model and the non-diabetic model in Fig. 7 had contributions from both diabetic cross-bridges and diabetic Ca^2+^ handling (Fig. 10B-G). The diabetic Ca^2+^ handling had a stronger effect on decreasing work than the diabetic cross-bridge (Fig. 10B). Conversely, the diabetic cross-bridges had a larger effect on the decreased shortening velocity (Fig. 10E), shortening power (Fig. 10F) and cross-bridge energy consumption (Fig. 10D). The increased cross-bridge efficiency in the diabetic model was due to diabetic cross-bridges rather than diabetic Ca^2+^ handling, as evidenced by the similarity of the two model simulations with diabetic cross-bridges (unfilled orange squares and unfilled purple circles; Fig. 10G).

#### 3.5.2 Parameter sensitivity analysis

The nature and degree of influence of each model parameter on the simulation outputs were explored in a systematic parameter sensitivity analysis in Fig. 11. The most striking result is that most parameters influence nearly all simulation output variables, emphasising the complexity of the interactions between the processes we are modelling. On average, Ca_50_ and Ca^2+^ twitch duration (*t*_95_) are the two parameters that have the greatest influence on the outputs; they are the most influential parameter for five of the outputs (systolic stress, maximum power, twitch duration, maximum energy consumption and maximum efficiency). Cross-bridge efficiency is the most sensitive output, as it is influenced by the majority of the parameters. In particular, lower values of 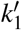 and 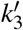, higher *k*_dATP_ and lower passive parameters can all contribute to greater cross-bridge efficiency. These changes are consistent with the effect of diabetes and provide a mechanistic explanation for the increase in cross-bridge efficiency predicted by the cross-bridge model (Fig. 7F).

**Figure 11:**
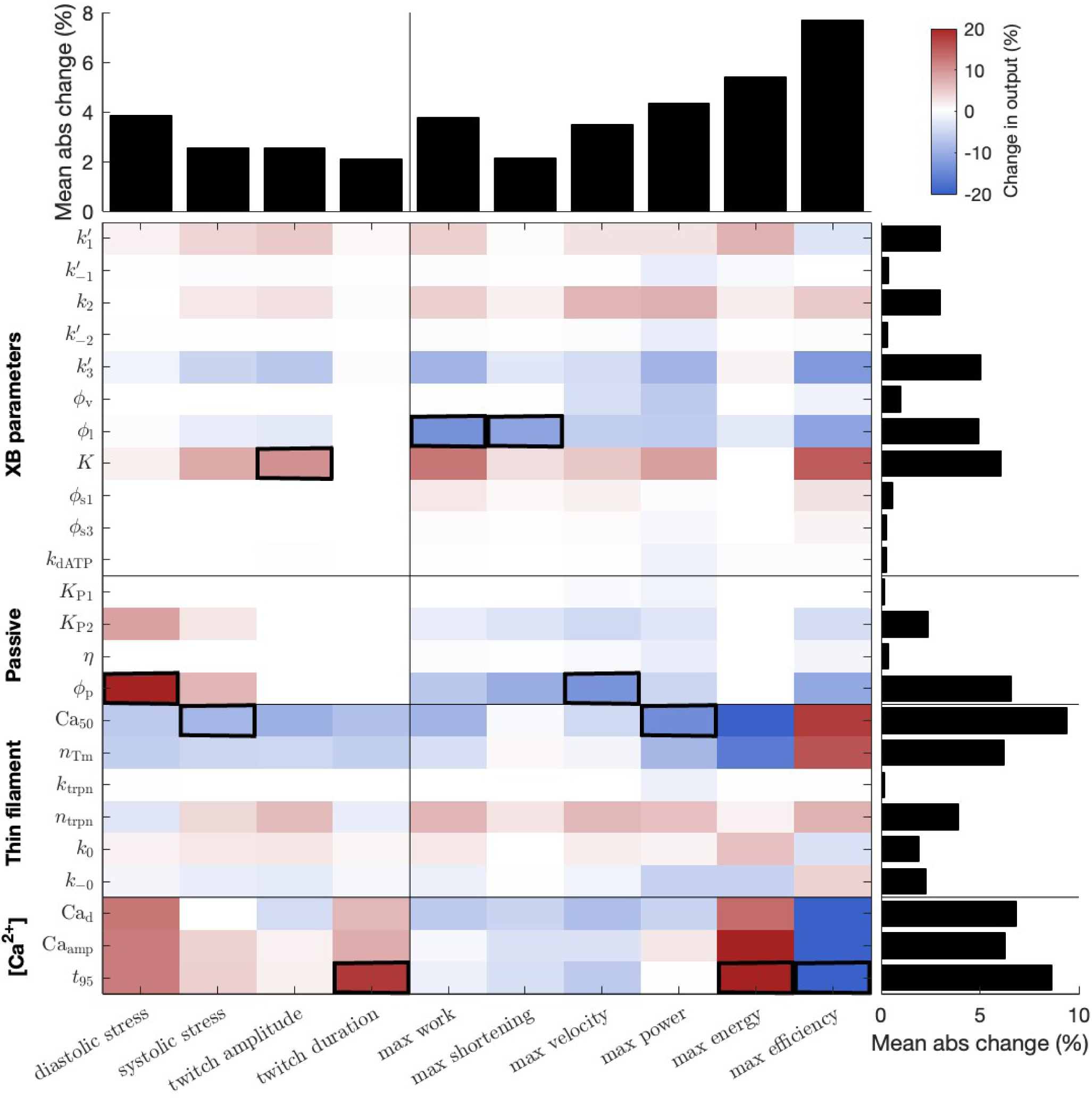
Sensitivity analysis of all parameters in the non-diabetic muscle model. Each parameter (rows) was increased by 10% and the consequent change in 10 output variables (columns) was recorded. The left-side y-axis lists the individual parameters, while the bottom x-axis lists the simulation outputs of interest. The bar plot along the top quantifies the sensitivity of each output, while the bar plot along the right side quantifies the influence of each input parameter. Squares with thick borders represent the parameter with the greatest absolute influence on an output variable.

## 4 Discussion

In this study, we have developed two models of human cardiac muscle function, one representing non-diabetic muscles and the other representing diabetic muscles. To our acknowledge, these are the first set of cross-bridge models that have been parameterised using data from human atrial tissues. The model simulations and subsequent analyses deliver deeper insights into the impact of diabetes on cardiac muscle mechanoenergetics. We discuss these findings in detail below, focussing on insights into the development of diabetic cardiomyopathy. These insights are drawn from both the parameterisation of the non-diabetic and diabetic cross-bridge models and the simulation of intact muscle behaviour, where the models were coupled to Ca^2+^ handling dynamics.

### 4.1 Insights into diabetic cardiac function from the cross-bridge model

In Musgrave et al. [13], measurements from maximally-activated human atrial trabeculae revealed several key differences between non-diabetic and diabetic muscles. These included lower stress generation, lower cross-bridge stiffness, slower cross-bridge cycling and enhanced sensitivity to ATP in diabetic muscles. In this study, parameterisation of cross-bridge models using these data (complex modulus and steady-state stresses) provided further support for the proposed mechanisms underlying these differences.

The optimisation of cross-bridge model parameters highlighted the differences in cross-bridge cycling rates between the non-diabetic and diabetic groups. The leftward shift measured in the diabetic complex moduli and the reduced proportion of the *α*-myosin isoform suggested slower cycling rates in diabetic muscle [13]. On the whole, this was reflected in the cross-bridge model parameterisation, where, under baseline conditions, all but one of the cycling rates were lower in the diabetic model (Table 3). Crucially, the values of both *k*_−1_ and *k*_3_ were lower in the diabetic model, indicating slower detachment rates. This, in turn, leads to lower cut-off frequencies for the high-pass transfer functions which make up the complex modulus response [14], explaining the leftward shift seen experimentally.

A key experimental finding of Musgrave et al. [13] was that diabetic muscles exhibited lower steady-state stresses and active stiffnesses. However, this difference was eliminated when the myofilament fraction (proportion consisting of only myocytes) within the muscles was taken into account. Considering the relevant parameters in the cross-bridge models reveals another level of complexity underlying this phenomenon. The value of *K*, the stiffness coefficient, is higher in the non-diabetic model, indicating that there is either a higher density of cross-bridges or a greater individual cross-bridge stiffness. Based on the structural information obtained from these muscles, we decomposed *K* into components representing the myofilament density (*ρ*_myo_) and the stiffness coefficient of cross-bridges (*K*_myo_; Eq. 2). Under this decomposition, the stiffness coefficient of cross-bridges was still appreciably higher in the non-diabetic model (Table 3). However, this still results in similar steady-state stresses in the myofilament-normalised cross-bridge models because the proportion of attached cross-bridges was higher in the diabetic model, a consequence of the slower detachment rates discussed above. A lower value of *K*_myo_ in the diabetic model means that there is either a lower density of cross-bridges within the myocytes of these muscles or the cross-bridges themselves are less stiff. One or both of these factors must be true in order to reconcile the similar stresses measured when normalising to myofilament fraction. Quantifying the myofilament fraction at the myocyte level using high-resolution electron microscopy (e.g. Rajagopal et al. [25]) could provide additional evidence to identify the source of this difference.

Model parameter fitting also provided insights into the differential responses of diabetic muscles to ATP that were observed in the experimental measurements. Since ATP is incorporated using a rapid equilibrium binding mechanism in the cross-bridge models, a *k*_d_ value was fitted to represent the sensitivity of the models to ATP concentration. In the non-diabetic model, *k*_dATP_ was 0.281 mM, which is higher than the half-activation value for ATP previously measured in rat cardiac trabeculae [26], but falls within typical ranges for other muscle fibres [27, 28]. In contrast, the diabetic model yielded an optimal *k*_dATP_ of 5 mM, the maximum allowed by the optimisation. Although this is well outside the expected range, it aligns with the unusual ATP sensitivity observed experimentally in Musgrave et al. [13]. This finding suggests that a heightened sensitivity to ATP in diabetic tissues underlies the difference in response.

### 4.3 Insights into diabetic cardiomyopathy development from muscle model simulations

While analysis of the non-diabetic and diabetic cross-bridge models allowed identification of the mechanisms driving differences in behaviour at maximal Ca^2+^ activation, simulations of whole muscle behaviour, which incorporated dynamic Ca^2+^ handling, provided a more physiologically relevant context for these findings.

#### Isometric twitches

In line with the lower steady-state stresses observed in the cross-bridge model, whole-muscle simulations predicted a lower isometric twitch amplitude in the diabetic model (Fig. 5A). When simulations accounted for the myofilament fraction to calculate myocyte stress, the difference in twitch amplitude was diminished (Fig. 5B). However, unlike the findings in Musgrave et al. [13], where normalising steady-state stresses with myofilament fraction eliminated the difference, the disparity in twitch amplitude persisted. This suggests that diabetes impacts twitch amplitude through a separate mechanism. Our simulations in Fig. 10A reveal that altered Ca^2+^ handling dynamics in diabetes also contribute to the decrease in twitch amplitude. The decrease arises from a combination of reduced Ca^2+^ transient amplitude (Fig. 3) and reduced Ca^2+^ sensitivity (Table 1) in diabetes.

The isometric contraction simulations in Fig. 5A support the hypothesis proposed by Musgrave et al. [13] that slowed cross-bridge kinetics in diabetes contributes to diastolic dysfunction. These slower rate constants led to a twitch duration that is 15 % longer in the diabetic model in simulations where the Ca^2+^ transients are identical (Fig. 10A). This finding aligns with experimental data on isometric twitch duration in both rat ventricular trabeculae [10] and human atrial trabeculae [11]. Although the simulated muscles fully relaxed under 1 Hz stimulation, it is plausible to speculate that the diabetic model might struggle to relax completely at higher frequencies.

#### Work-loop mechanics

The muscle model also enabled predictions of the behaviour of non-diabetic and diabetic muscles under stress-length work-loops, which more closely mimic the loading conditions experienced by tissues in the heart [22]. Key differences revealed in the simulations were that, compared to the non-diabetic model, the diabetic model had: lower work done, lower maximum shortening velocity, much lower power of shortening, much lower cross-bridge enthalpy and higher cross-bridge efficiency (Fig. 7). These trends appear to primarily reflect differences in cross-bridge function rather than differences in Ca^2+^ handling, as most differences persisted even when the diabetic model was paired with non-diabetic Ca^2+^ dynamics (Fig. 10, purple unfilled circles). Additionally, the trends were largely independent of differences in muscle passive force, as they remained evident in simulations where both models used the same non-diabetic passive parameters (Fig. 10). For all the work-loop mechanical outputs, increasing the passive parameters (i.e. increasing stiffness) in the parameter sensitivity analysis led to a reduction in their magnitudes (Fig. 11). Consequently, if the diabetic passive properties were stiffer, the differences from the non-diabetic model would be further amplified for all of these outputs. This is particularly relevant to note as the findings of lower passive stress in Musgrave et al. [13] are somewhat contradictory with the diastolic dysfunction commonly seen in diabetic cardiomyopathy [2].

The effect of diabetes on the magnitude of the shortening velocity aligns well with maximum unloaded shortening rates measured in type 1 diabetic rat papillary muscles [29]. Prolonged shortening durations have also been reported previously in type 2 diabetic rats [30]. In our diabetic model, the maximum shortening velocity is primarily due to a reduced attachment rate, *k*_1_ (Fig. 11). Shortening power is the product of this velocity and afterload and therefore reflects two work-loop outputs that are lower in the diabetic model.

The reduced work and power predicted by the diabetic muscle model suggest systolic dysfunction that could extend to the organ level. In particular, shortening power is less sensitive to changes in *K* compared to work (Fig. 11). As a result, shortening power would still be markedly reduced in the diabetic muscle, even if the myofilament density was the same between groups.

#### Work-loop energetics

The reduced energy consumption per twitch (change in enthalpy) in the diabetic model (Fig. 7C) reflects the slower rate of cross-bridge detachment, *k*_3_, and, to a lesser extent, the lower density of myofilaments (Eq. 24). Since the difference in work done by both models was less pronounced than the difference in the change in enthalpy, this led to higher efficiency in the diabetic muscle: a given unit of work being done for much less energy consumption. The parameter sensitivity analysis (Fig. 11) revealed that cross-bridge efficiency is determined by multiple parameters in the model. In diabetes, the higher cross-bridge efficiency was driven by slower attachment and detachment rates (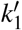 and 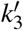), higher *k*_dATP_ (which further lowers *k*_3_ under baseline conditions) and lower passive model parameters (especially *K*_P2_ and *ϕ*_p_) compared to the non-diabetic model.

The values for the cross-bridge efficiency predicted by the model simulations (Fig. 7F) were two to three times lower than those previously measured in rat ventricular trabeculae [10, 31]. This discrepancy could be attributed to species and chamber differences, the advanced age of the human patients from whom the cross-bridge data were derived, or uncertainties in the enthalpy calculations. Specifically, the density of cross-bridges in a muscle (*ρ*_xb_ in Eq. 24) is challenging to estimate precisely, so the relative differences between the two groups are the most relevant metric for enthalpy and efficiency outputs. While we predicted considerable differences in the cross-bridge efficiency between the two groups, peak mechanical efficiency measured from cardiac muscles of type 1 diabetic rats has not been shown to be different compared to controls [10, 29]. Cross-bridge efficiency compares the work done by the muscle to the energy consumed by cross-bridge cycling, whereas mechanical efficiency compares the work done to the energy consumed by all processes in the muscle.

To predict the mechanical efficiency using our muscle models, other major sources of energy consumption in the myocyte, such as the sarcoendoplasmic reticulum Ca^2+^ATPase (SERCA) and the Na^+^/K^+^ pumps, would need to be considered. Increased SERCA channel density has previously been reported in diabetic human atrial tissue [11]and is thought to be a compensatory mechanism for impaired relaxation in diabetes. Combined with findings of elevated Ca^2+^ leak from the sarcoplasmic reticulum by Jones [32], there is strong evidence that SERCA may consume more energy in diabetic tissues, potentially resulting in comparatively lower mechanical efficiency.

In Musgrave et al. [13], a decrease in the proportion of *α*-myosin isoform was identified as a likely mechanism underlying the observed slower cross-bridge cycling rates. This hypothesis is supported by findings of reduced *α*-myosin proportions in diabetic rat hearts [33, 34], and in the ventricles of failing human hearts [35]. Since the *α*-myosin isoform is associated with higher shortening velocities and greater power generation at the expense of higher energy consumption [36], a lower proportion of this isoform aligns with the predominant effects of diabetes on work-loop outputs in our simulations.

#### Metabolite sensitivity in the muscle model

Given that diabetic cardiomyopathy is associated with metabolic dysfunction, the effects of altered metabolites were investigated. The lower value of *k*_−1_ found in the diabetic model reflects a lower sensitivity to P_i_. This manifested as a less pronounced response to increases in P_i_ concentration compared to the non-diabetic model (Fig. 8). As raised P_i_ generally causes negative responses (lower stress produced, lower work done and lower power of shortening), this could be considered a compensatory mechanism which protects the diabetic muscle from the effects of increased P_i_ in the *in vivo* heart [7]. However, while lower sensitivity means that the diabetic muscle is less affected by elevated P_i_, the systolic dysfunction observed under the baseline metabolic conditions will still worsen as P_i_ concentration rises.

Conversely, the high value of *k*_dATP_ found in the diabetic model reflects a much greater sensitivity to physiological concentrations of ATP, consistent with the experimental findings in Musgrave et al. [13]. When simulating the effect of lowered ATP, which may occur in diabetic muscle *in vivo* [6], a mix of responses was observed (Fig. 9). On the positive side, lower ATP resulted in less energy consumed which increased cross-bridge efficiency. There was also an increase in systolic stress and work done, the latter being much more pronounced in the diabetic model due to its heightened sensitivity to ATP. However, the diabetic model also exhibited significant increases in diastolic stress and twitch duration, along with a decrease in shortening velocity under reduced ATP conditions. These findings suggest that increased sensitivity to ATP and lower ATP concentration in diabetic muscle could contribute to the diastolic dysfunction characteristic of diabetic cardiomyopathy.

#### Insights into treating diabetic cardiomyopathy

Based solely on the experimental data used to inform the model simulations, it was clear that both cross-bridge function and Ca^2+^ handling exhibited signs of impairment in diabetic hearts [9, 13]. To better understand the individual contributions of these two components to the overall diabetic muscle response, we designed a set of cross-over simulations to isolate their effects (Fig. 10). These simulations also provide a framework for evaluating the potential impact of treatments aimed at restoring each of these target areas to non-diabetic levels.

The deficiencies of the diabetic model could not be fully corrected by replacing either the Ca^2+^ handling or the cross-bridge components with their non-diabetic counterparts. However, restoring the cross-bridge properties led to greater improvements in twitch duration, shortening velocity, and shortening power, while restoring Ca^2+^ handling resulted in greater enhancements in twitch amplitude and the work done by the muscle (Fig. 10).

Since the differences in diabetic Ca^2+^ handling dynamics involve elevated diastolic Ca^2+^ levels and slower decay rates, targeting SERCA may be able to reverse these changes. As for the cross-bridge function, the emerging use of myosin-modulating drugs to target heart failure and cardiomyopathies seems promising [37]. However, it is important to recognise that the two key issues - reduced stress development and slower cross-bridge cycling-present conflicting objectives for such drugs. Myosin activators like omecamtiv and danicamtiv enhance force production but at the cost of slower cross-bridge cycling rates [38]. Conversely, myosin inhibitors like mavacamten and aficamten can increase cross-bridge cycling rates (which may be particularly important under conditions of lower ATP availability) but reduce contractility [37]. Nevertheless, the models presented here provide a foundation for exploring the potential impact of these drugs in treating diabetic heart disease.

### 4.3 Model considerations

In this study, our model predictions extended beyond the permeabilised, maximally-activated conditions under which the experimental data were collected. It has been well-established that experimental measurements acquired from permeabilised muscles are not always equivalent to their intact counterparts [39]. In a multiscale modelling study of the human heart, Land et al. [20] found that the cross-bridge model they had parameterised using permeabilised experimental data could not produce sufficient force or shortening to adequately perform work-loops at the organ level. While we have not extended the model to the organ scale, the stress-length work-loops and isometric contractions performed by our muscle models suggest that the parameters identified from permeabilised muscle measurements represent intact trabeculae well. In particular, the peak stresses obtained from isometric contraction simulations of non-diabetic and diabetic muscle (Fig. 5) fall within the standard error of those obtained in similar intact human trabeculae [9]. This model output is determined primarily by the steady-state stresses obtained experimentally, which are known to vary between intact and permeabilised preparations [39]. The velocities of shortening obtained during the work-loops (Fig. 7) are typical of those obtained in intact rat trabeculae [23, 31]. This output is characterised by the frequency information contained in the complex modulus and is typically not considered to be affected by muscle permeabilisation [40]. In contrast, the Land et al. [20] study used rapid length step experiments to parameterise their cross-bridge kinetics, which later required adjustments for integrating within a whole organ model. This suggests that the complex modulus measurement provides a more reliable approach for deriving cross-bridge kinetic information. Due to the very small length changes involved in this protocol, it is feasible that the muscles are also less sensitive to the length-related discrepancies typically observed in permeabilised muscles [39] when performing these experiments.

Musgrave et al. [14] proposed that fitting human complex modulus data, as opposed to rat data, would require a different combination of metabolite and strain dependencies. In the present study, while strain dependence on *k*_3_ was retained from the rat model, an additional strain dependence on a rate unaffected by ATP or P_i_ was also required. Through optimisation, *k*_1_ was identified as the most suitable option. This reflects a reduced probability of cross-bridge binding when there is positive strain in state **B**. Another departure from the rat model [14] involved the mechanism for P_i_ binding. Although rapid equilibrium binding for P_i_ has been supported by evidence [14, 41, 42], its more complex effects led to poorer fits for both non-diabetic and diabetic models, and it was consequently excluded from the final human model. Additionally, to align with current theoretical frameworks of understanding [43, 44], we explored incorporating P_i_ directly at the reverse power stroke step *k*_−2_, rather than the unbinding step *k*_−1_. However, this approach produced a significantly worse fit to the diabetic data and was hence rejected.

Another difference identified when fitting the models to the human data was the potential existence of another process in the complex modulus at low frequencies. Kawai et al. [45] initially found that the low frequency ‘A’ process seen in skeletal muscle is absent in cardiac muscles. However, subsequent studies revealed that it can manifest in cardiac muscle at higher temperatures [46], which fall within the range of the experimental data used in this study. Despite this, none of the mechanisms typically seen in biophysical cross-bridge models [17] are capable of reproducing this behaviour. This process reflects a much slower response (< 0.5 Hz) compared to other processes, and its absence is unlikely to materially impact the findings of this study.

### 4.4 Conclusions

In this study, we have developed metabolite-sensitive models of human atrial cross-bridge and muscle function for both non-diabetic and diabetic conditions.

The parameterisation of these cross-bridge models offered mechanistic insights into two key findings in diabetic human atria data. The leftward shift in the complex modulus of diabetic tissues was attributed to slower cross-bridge detachment, while the reduced active stress development resulted from a decrease in the stiffness of cross-bridges in diabetic muscle.

When simulating physiological contractions using the muscle models, these differences in cross-bridge cycling rates and cross-bridge stiffness led to further differences in function between the non-diabetic and diabetic groups. The diabetic model produced isometric twitches that were lower in amplitude and slower in time course, and work-loop contractions that shortened more slowly, produced less work and developed lower power. However, due to its slower detachment rates, the diabetic model consumed much less ATP, resulting in greater cross-bridge efficiency. While altered Ca^2+^ handling in diabetes also contributed to some of these outcomes, cross-bridge function was identified as the primary driver of most observed differences.

Simulating potential metabolic dysfunction provided additional insights into the development of diabetic cardiomyopathy. The diabetic model’s lower sensitivity to P_i_ suggested a compensatory mechanism that may help mitigate the adverse effects of elevated P_i_ concentrations in the diabetic heart. On the other hand, reduced ATP concentration appeared to contribute to diastolic dysfunction in diabetic muscles. The diabetic model exhibited heightened sensitivity to changes in ATP, leading to prolonged twitch duration and elevated diastolic stress levels under conditions of reduced ATP. These findings highlight the critical role of metabolic alterations in the patho-physiology of diabetic cardiomyopathy.

## Additional Information

## Data availability statement

Complete model code to reproduce all simulation figures is available at https://github.com/JuliaMusgrave/AtrialModel_2025_Human.

## Author Information

JM developed the model, performed simulations and drafted the manuscript. All authors contributed to the conception of the work, critically revised the manuscript and approved the final version.

## Appendix - complete model description

### Muscle model equations

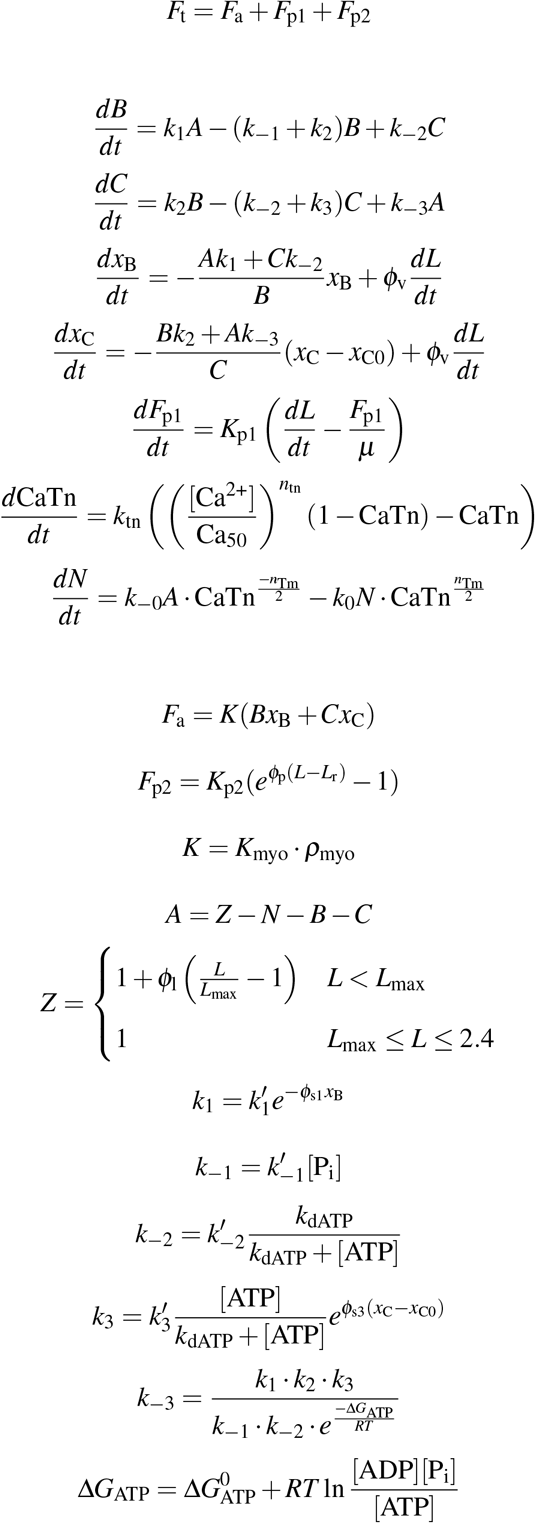

**Table 4:**
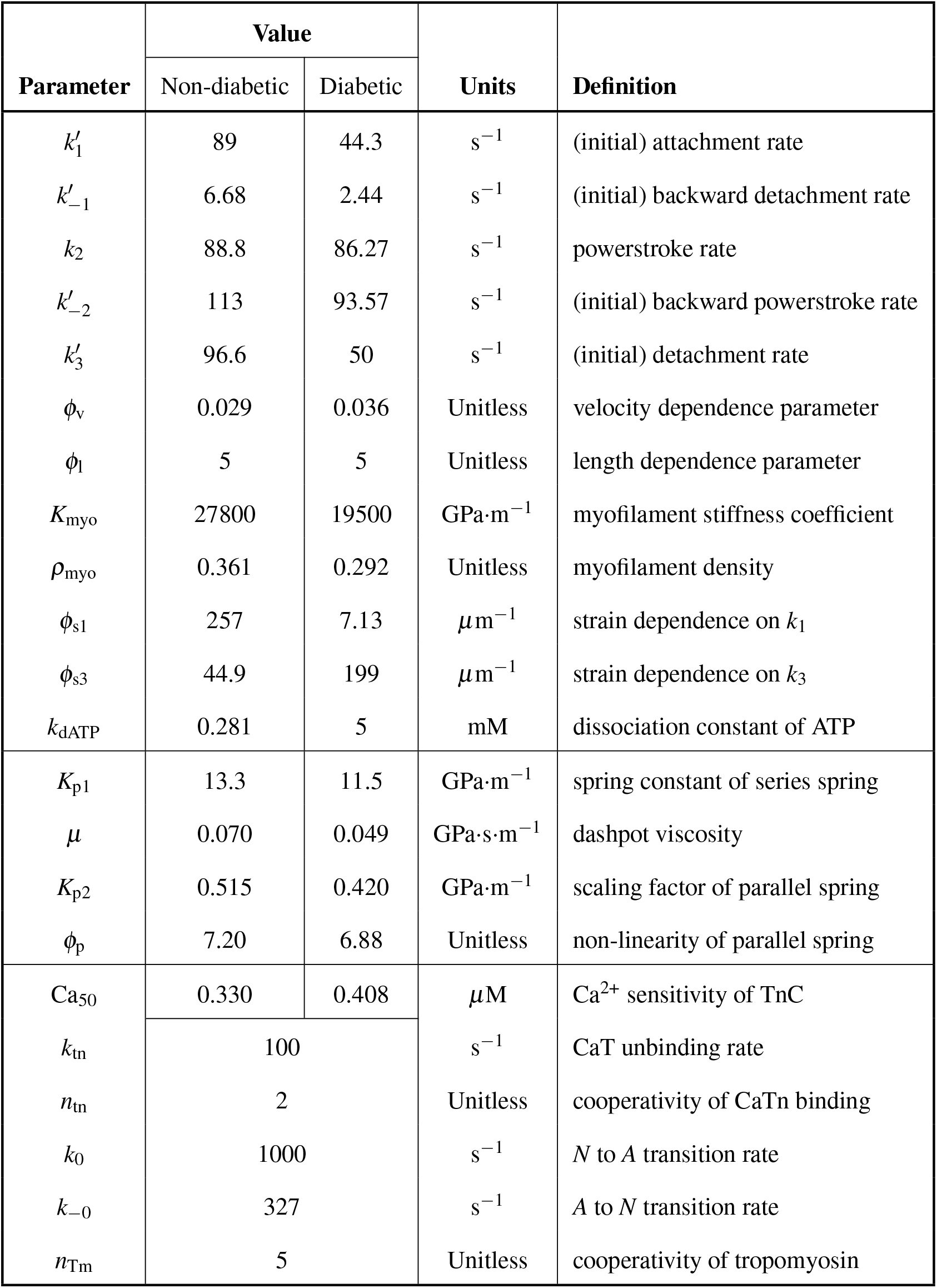
List of parameters for the non-diabetic and diabetic versions of the human muscle model. The cross-bridge and passive parameters come from parameterisation of the model to non-diabetic and diabetic human atrial trabeculae data from Musgrave et al. [13]. The thin filament regulation parameters are taken from Land et al. [20], except for Ca^2+^ sensitivity which is taken from Jones et al. [9].

| Parameter | Value |  | Units | Definition |
| --- | --- | --- | --- | --- |
|  | Non-diabetic | Diabetic |  |  |
| $k'_1$ | 89 | 44.3 | $s^{-1}$ | (initial) attachment rate |
| $k'_{-1}$ | 6.68 | 2.44 | $s^{-1}$ | (initial) backward detachment rate |
| $k_2$ | 88.8 | 86.27 | $s^{-1}$ | powerstroke rate |
| $k'_{-2}$ | 113 | 93.57 | $s^{-1}$ | (initial) backward powerstroke rate |
| $k'_3$ | 96.6 | 50 | $s^{-1}$ | (initial) detachment rate |
| $\phi_v$ | 0.029 | 0.036 | Unitless | velocity dependence parameter |
| $\phi_l$ | 5 | 5 | Unitless | length dependence parameter |
| $K_{myo}$ | 27800 | 19500 | $GPa \cdot m^{-1}$ | myofilament stiffness coefficient |
| $\rho_{myo}$ | 0.361 | 0.292 | Unitless | myofilament density |
| $\phi_{s1}$ | 257 | 7.13 | $\mu m^{-1}$ | strain dependence on $k_1$ |
| $\phi_{s3}$ | 44.9 | 199 | $\mu m^{-1}$ | strain dependence on $k_3$ |
| $k_{dATP}$ | 0.281 | 5 | mM | dissociation constant of ATP |
| $K_{p1}$ | 13.3 | 11.5 | $GPa \cdot m^{-1}$ | spring constant of series spring |
| $\mu$ | 0.070 | 0.049 | $GPa \cdot s \cdot m^{-1}$ | dashpot viscosity |
| $K_{p2}$ | 0.515 | 0.420 | $GPa \cdot m^{-1}$ | scaling factor of parallel spring |
| $\phi_p$ | 7.20 | 6.88 | Unitless | non-linearity of parallel spring |
| $Ca_{50}$ | 0.330 | 0.408 | $\mu M$ | $Ca^{2+}$ sensitivity of TnC |
| $k_{tn}$ | 100 | | $s^{-1}$ | CaT unbinding rate |
| $n_{tn}$ | 2 | | Unitless | cooperativity of CaTn binding |
| $k_0$ | 1000 | | $s^{-1}$ | $N$ to $A$ transition rate |
| $k_{-0}$ | 327 | | $s^{-1}$ | $A$ to $N$ transition rate |
| $n_{Tm}$ | 5 | | Unitless | cooperativity of tropomyosin |

**Table 5:** List of variables and constants for the human muscle model.

| Parameter | Value | Units | Definition |
| --- | --- | --- | --- |
| $F_t$ | output variable | kPa | total muscle stress |
| $B$ | state variable | Unitless | proportion of XBs in state **B** |
| $C$ | state variable | Unitless | proportion of XBs in state **C** |
| $x_B$ | state variable | $\mu\text{m}$ | strain in state **B** |
| $x_C$ | state variable | $\mu\text{m}$ | strain in state **C** |
| $F_{p1}$ | state variable | kPa | passive stress from spring/dashpot |
| CaTn | state variable | Unitless | proportion of TnC with $\text{Ca}^{2+}$ bound |
| $N$ | state variable | Unitless | proportion of non-permissible XBs |
| $L$ | input variable | $\mu\text{m}$ | sarcomere length |
| $[\text{Ca}^{2+}]$ | input variable | $\mu\text{M}$ | concentration of intracellular $\text{Ca}^{2+}$ |
| $[\text{P}_i]$ | input variable | mM | concentration of $\text{P}_i$ |
| $[\text{ATP}]$ | input variable | m | concentration of ATP |
| $F_a$ | algebraic variable | kPa | active stress |
| $F_{p2}$ | algebraic variable | kPa | passive stress from parallel spring |
| $K$ | algebraic variable | $\text{GPa}\cdot\text{m}^{-1}$ | muscle XB stiffness coefficient |
| $A$ | algebraic variable | Unitless | proportion of XBs in state **A** |
| $Z$ | algebraic variable | Unitless | proportion of XBs in binding range |
| $k_{-3}$ | algebraic variable | $\text{s}^{-1}$ | backward attachment rate |
| $\Delta G_{\text{ATP}}$ | algebraic variable | $\text{kJ}\cdot\text{mol}^{-1}$ | Gibbs free energy of ATP |
| $L_{\text{max}}$ | 2.3 | $\mu\text{m}$ | maximum sarcomere length |
| $x_{C0}$ | 0.01 | $\mu\text{m}$ | powerstroke size |
| $\Delta G_{\text{ATP}0}$ | -30 | $\text{kJ}\cdot\text{mol}^{-1}$ | standard Gibbs free energy of ATP |
| $R$ | 8.314 | $\text{J}\cdot\text{mol}^{-1}\cdot\text{K}^{-1}$ | universal gas constant |
| $T$ | 310 | K | temperature |
| $[\text{ADP}]$ | 0.036 | mM | concentration of ADP |
| $t$ | independent variable | s | time |

### Additional equations for work-loop contraction simulations

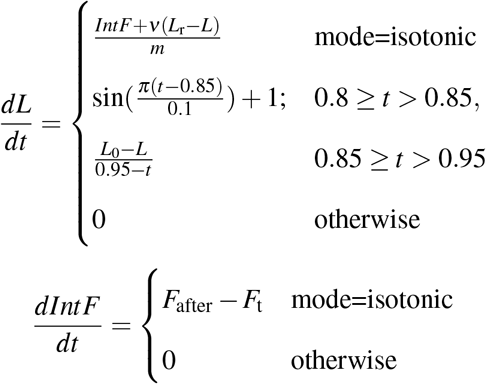

**Table 6:** List of variables and constants for running work-loop simulations in the human muscle model.

| Parameter | Value | Units | Definition |
| --- | --- | --- | --- |
| $IntF$ | state variable | kPa·s | integral of force |
| $F_{\text{after}}$ | input variable | kPa | afterload stress |
| $v$ | 0.00005 | GPa·s·m<sup>-1</sup> | viscosity |
| $m$ | 0.00001 | GPa·s<sup>2</sup>·m<sup>-1</sup> | mass |
| $L_0$ | 2.2 | μm | starting muscle length |

